# Molecular logic of pheromone recognition enables rational design of insect behavior modulators

**DOI:** 10.1101/2025.05.30.657074

**Authors:** Sukjin S. Jang, Sasha A. T. McDowell, Pramod KC, Marija Simjanoska, Sanjana Mandala, Xiao Zhang, Hanjie Jiang, Philip A. Cole, Jeffrey Riffell, Qinheng Zheng, Josefina del Mármol

## Abstract

Reproductive success across animal taxa relies on the unequivocal identification of volatile chemical cues used for sexual communication. These chemical signals, called pheromones, are often structurally nearly identical, sometimes differing only by the oxidation state of a functional group, yet elicit vastly different behaviors. The structural mechanisms driving this exquisite selectivity remain elusive, hindering the rational development of tools to modulate these pathways. Here, we decode the structural logic of pheromone recognition to rationally manipulate insect behavior. Using cryo-electron microscopy and functional mutagenesis, we show that the silkmoth Bombyx mori distinguishes between two highly similar compounds through the selective formation of a thio-hemiaminal, an uncommon covalent bond between the receptor and the pheromone. We leveraged these structural insights to undertake the first rational design of small molecules to disrupt insect mating. By engineering targeted electrophiles that irreversibly anchor to this reactive pocket, we successfully abolished long-range mate-tracking behavior in an agricultural pest model in vivo. This work identifies an unrecognized chemical strategy driving olfactory specificity and provides a blueprint for the rational design of airborne behavior modulators.

## Introduction

From triggering suckling behavior in newborn mammals to guiding male moths in long-range pursuit of females, volatile chemical cues are the medium of critical social information in the animal kingdom [1–3]. A key unresolved challenge is understanding how olfactory systems accurately sense these chemical cues termed ‘pheromones’, which often differ from one another by as little as a single atom and yet elicit fundamentally different physiological and behavioral responses [3–5]. High chemical specificity is a hallmark of pheromone communication to ensure that innate responses mediating threat avoidance, kin recognition or mating are triggered only by the correct signal [2, 6, 7].

A structural understanding of how these volatile chemical cues are sensed will not only illuminate the mechanisms of sexual communication but would also enable the rational development of highly specific chemical tools to modulate animal behavior and re-engineer ecological interactions. In particular, targeted control of insect mating holds great potential [1, 6, 8]. Insects, comprising over 50% of all animal species on earth [9], include some of the most destructive invasive pests and disease vectors, ravaging crops and transmitting deadly diseases that account for billions of dollars in agricultural losses and almost one million deaths every year [10–13]. Because of their profound impact on human life, insect pheromone sensing pathways represent attractive targets for chemical intervention with the potential to reshape global health and agriculture [8]. However, rational design of synthetic compounds to manipulate insect mating requires elucidation of the underlying structural principles that govern pheromone communication in insects.

The term pheromone was first coined to describe the ‘invisible’ mating signals of moths [14]. Female moths release a species-specific pheromone blend alerting males of reproductive receptivity. Males must detect and identify the chemical signature of the pheromone blend and track its plume over vast distances. Because these insects often exist at low population densities and mate at night, reproductive success depends on male pheromone receptors that can unequivocally discriminate amongst trace amounts of nearly identical pheromones. This sensory prowess is as impressive as it is destructive: moth species pose great danger to the global food supply, causing the loss of 10 to 16% of major grain crops and costing the global economy billions of dollars [10, 15, 16].

Here, we define the structural basis of sex communication in moths and uncover a previously unrecognized chemical strategy for specific pheromone recognition. Using structural and biochemical analyses, we show that aldehyde pheromones form an uncommon, reversible covalent bond with their receptors. This specific chemical interaction enables unequivocal differentiation from other highly similar volatile ligands. Leveraging this structural information, we designed ‘mating modulators’, synthetic compounds that irreversibly target pheromone receptors with high specificity and efficacy to disrupt insect behavior. Lastly, we show that these rationally designed compounds are effective in modulating mating behavior *in vivo*. These findings uncover new structural mechanisms of chemical specificity in the olfactory system and provide a framework for the structure-based design of chemical tools to modulate animal behavior for societal benefit.

### A modular structural strategy to detect the combinatorial chemistry of lepidopteran pheromones

We focused our study on the sex communication of lepidopterans (moths and butterflies), whose long-range pheromone communication system has been extensively studied because of their pervasive roles in causing agricultural and ecological damage [1, 6, 8]. Lepidopteran pheromones share a core chemical architecture: a long carbon chain with a variable number of unsaturations, capped by a functional group that is most frequently an aldehyde, an alcohol, or an acetate (Figure 1A) [1, 6, 17, 18]. Variations in chain length, number and stereochemistry of the double bonds, and identity of the functional group provide a rich combinatorial chemical canvas that allows more than 165,000 species of lepidopterans to evolve their own unique pheromone blend [1, 19].

**Figure 1.**
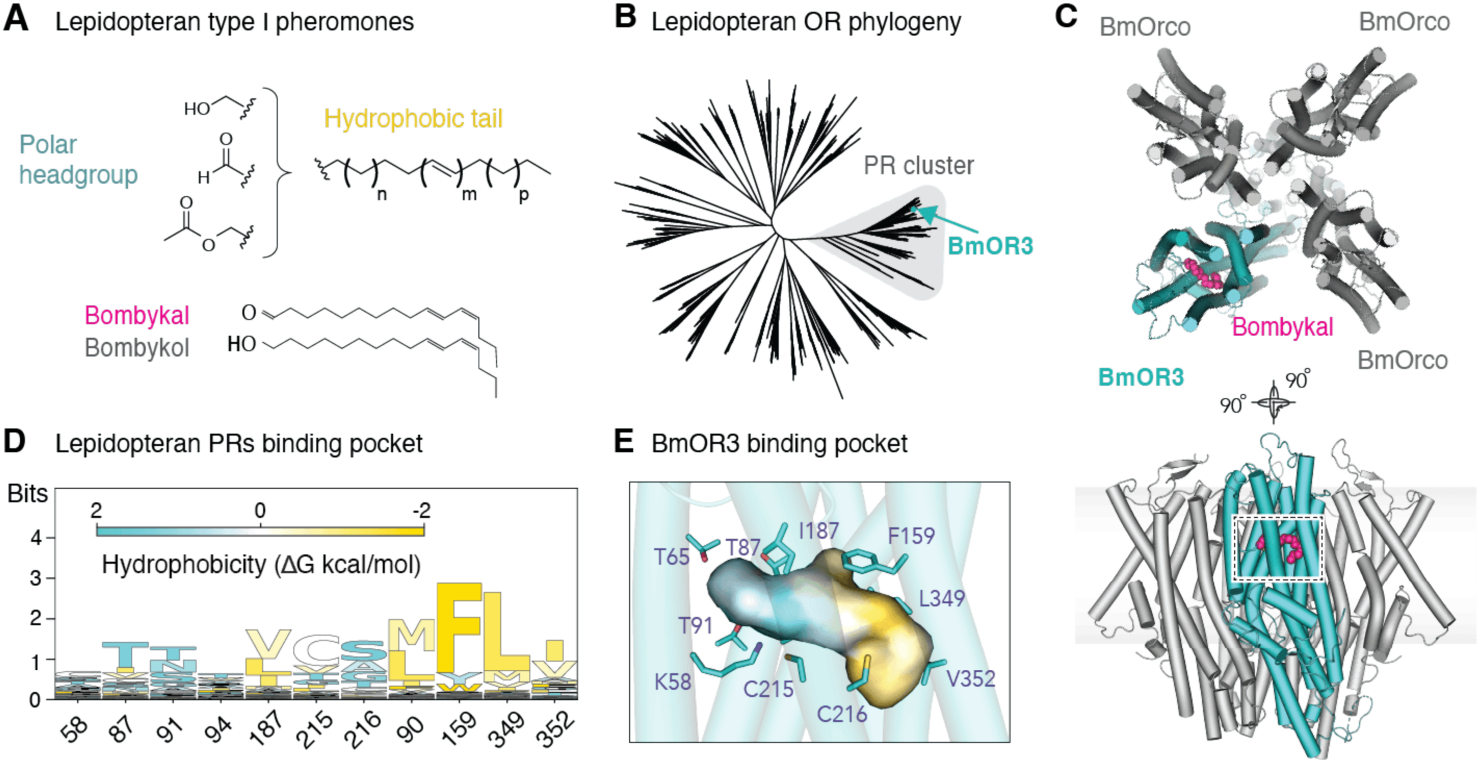
A conserved binding mode for pheromone recognition by lepidopteran PRs. (**A**) General chemical architecture of type I lepidopteran pheromones. A polar head group (typically an alcohol, aldehyde or acetate) is followed by an unbranched aliphatic tail with variable number of saturated (n, p) and unsaturated (m) carbons. Bombykal (magenta) and bombykol (grey) are shown below. (**B**) Unrooted phylogenetic tree of lepidopteran ORs highlighting the clustering of the PR subfamily (grey). Teal arrow indicates the position of BmOR3. (**C**) Top (upper) and side (lower) views of the cryo-EM structure of BmOR3/BmOrco. BmOR3 subunit is shown in teal, BmOrco subunits in grey, and bombykal in magenta. The grey band in the side view indicates the approximate position of the lipid membrane. (**D**) Sequence conservation of the PR binding pocket across 249 lepidopteran species. Letter height is proportional to its conservation at the indicated position. Residues are colored by hydrophobicity (gold, hydrophobic residues, aqua, polar residues) following the Wimley-White Octanol scale [47]. (**E**) View of the BmOR3/BmOrco binding pocket colored by hydrophobicity as in (**D**). Bombykal-interacting residues lining the binding pocket are annotated. The pocket surface was generated using CavitOmiX [48].

To detect these pheromones, lepidopteran males use pheromone receptors (PRs) located in their main olfactory organ, the antenna [4, 20–23]. PRs are a distinct cluster within the larger family of insect odorant receptors (ORs), a family of ligand-gated ion channels that initiate neuronal signaling upon binding odorants or pheromones (Figure 1B)[17, 24]. Lepidopteran pheromone receptors have so far eluded structural characterization, partly due to the biochemical complexity of the PR family and chemical complexity of their ligands, and therefore how this family evolved to decode the combinatorial chemistry of pheromonal signals is unexplored.

The subject of our study is the silkmoth *Bombyx mori*, a foundational model organism for olfaction research that presents the perfect opportunity for elucidating the basis of specificity in pheromone recognition. To attract males, *B. mori* females emit a pheromone blend consisting of a 10:1 ratio of bombykol and bombykal, two near-identical compounds that differ only in the oxidation state of their terminal head group (Figure 1A) [25]. Despite their highly similar chemical formulas, these compounds drive opposing behaviors: while trace amount of bombykol promote courtship and mating, excess amount of bombykal acts as a strong mating suppressor [25]. The male silkmoth olfactory system must therefore sense bombykal with high accuracy and unequivocally discriminate it from bombykol for mating success.

At the molecular level, bombykal recognition is primarily mediated by the PR BmOR3. Like most insect ORs, BmOR3 assembles with the obligate co-receptor Orco to form a hetero-tetrameric, calcium-permeable ligand-gated ion channel. To uncover the structural basis of pheromone sensing, we purified (Figure S1) and determined the cryo-EM structures of BmOR3/BmOrco both in the presence (2.68Å, Figure 1C, Figure S2A, S3A, Table S1) and absence (2.89Å, Figure S2B, S3B, Table S1) of bombykal. Consistent with recent studies of other OR/Orco complexes, BmOR3/BmOrco adopts a 1:3 stoichiometry [26–29]. All four subunits contribute to a central ion conduction pathway with a singular extracellular opening, a wide transmembrane vestibule and four intracellular exits (Figure S4A). The bombykal-bound structure exhibits an asymmetric widening of the channel pore driven by a diagonal displacement of the BmOR3 pore helix away from the main axis. This active conformation is consistent with structural observations of other ligand-bound insect OR/Orco complexes (Figure S4B-D)[26–29].

In the bombykal-bound structure, strong ligand density occupies a cavity ∼20Å deep in the BmOR3 transmembrane domain, formed by the convergence of helices S2, S3, S4, and S6 (Figure 1C and Figure S3A). While situated in the canonical insect OR binding site, the BmOR3 pocket is significantly larger and chemically segmented: the terminal region that packs against the S1 helix is lined with charged and polar residues, whereas the region extending towards the S6 helix forms a hydrophobic tunnel with a terminal curl (Figure 1D,E). In the ligand-free structure, a small density occupies only the polar end of the pocket, likely representing a small organic molecule (Figure S3B).

The bimodal architecture of the BmOR3 pocket mirrors the bimodal chemistry of the pheromone and suggests a modular mechanism for pheromone recognition, where polar residues coordinate the functional group and the hydrophobic tunnel dictates carbon tail geometry. To evaluate the extent of the conservation in this modular recognition strategy across the clade, we mapped the bombykal-interacting amino acids in the BmOR3 structure and evaluated their conservation across the lepidopteran PR family (Figure 1D). Indeed, sequence conservation follows a modular pattern. Residues lining the hydrophobic tunnel are largely conserved and always hydrophobic, whereas those coordinating the polar headgroup are highly variable and consistently polar. This analysis reveals an elegant evolutionary strategy by which lepidopteran PRs rely on a unified structural scaffold, where discrete pocket elements can be independently re-tuned to detect a highly diversified pheromone space, enabling rapid diversification of sexual communication signals.

### A covalent bond underlies chemical specificity of BmOR3 towards bombykal

To understand how BmOR3 selectively responds to bombykal and not bombykol, we first characterized the degree of specificity of BmOR3 *in vitro*. Using an established calcium flux assay in HEK293T, we found that BmOR3/BmOrco exhibits over ten-fold higher sensitivity for bombykal (EC_50_ = 0.100 ± 0.007 μM) over bombykol (EC_50_ = 1.49 ± 0.06 μM) (Figure 2A and Table S2). In addition, bombykol acts as a partial agonist, eliciting only ∼75% of the maximal bombykal response at saturating concentrations (Figure 2A and Table S2).

**Figure 2.**
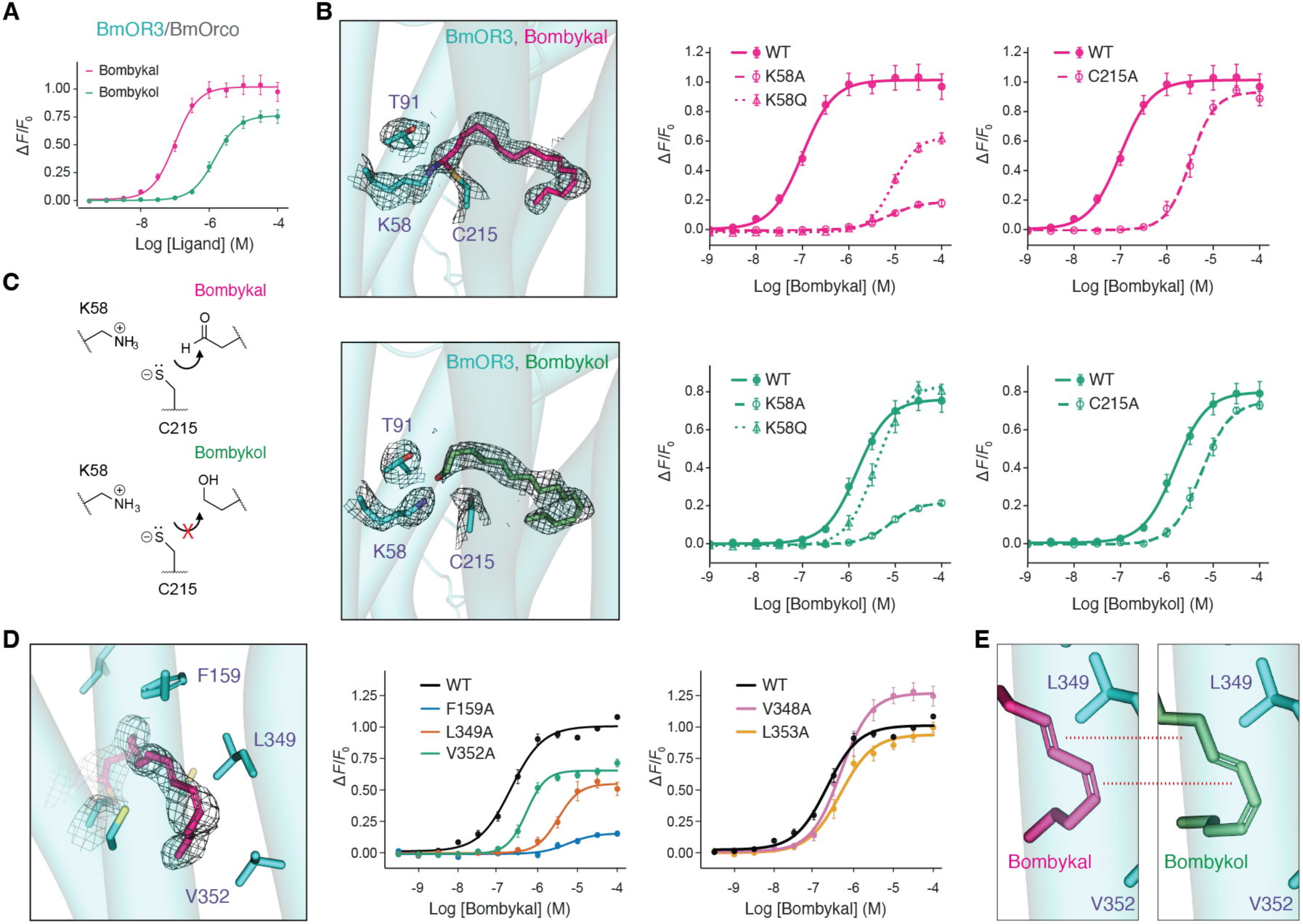
Structural mechanism of bombykal selectivity by BmOR3/BmOrco. (**A**) Dose-response curves of BmOR3/BmOrco to bombykal (magenta) and bombykol (green) in an in vitro cell-based calcium imaging assay. (**B**) Left, detail of the cryo-EM density of the BmOR3 binding pocket bound to bombykal (top) or bombykol (bottom). Right: Dose-response curves of Lys58 and Cys215 mutants to bombykal (top) and bombykol (bottom). (**C**) Schematics of the proposed chemical interaction between Lys58 and Cys215 and bombykal (top) or bombykol (bottom). (**D**) Left: Close-up view of the bombykal-bound pocket highlighting residues interacting with the hydrophobic tail of bombykal. Middle and right: Dose-response curves of alanine substitutions of residues contacting bombykal’s tail (middle) and neighboring residues that do not (right). (**E**) Close-up of the ligand-bound binding pocket showing the shifted position of the conjugated dienes of bombykal (left) and bombykol (right).

In our bombykal-bound structure of BmOR3, we observe that bombykal density extends continuously from Lys58 and Cys215, two reactive nucleophilic amino acids (Figure 2B and Figure S3A). Because the bombykal aldehyde is a strong electrophile (Figure 2C), this complementary reactivity prompted us to consider whether aldehyde recognition by BmOR3 might involve formation of a covalent bond. The identity of the chemical groups involved suggests formation of a thio-hemiaminal (Figure S5), a three-way covalent linkage connecting the thiol group of Cys215, the primary amine group of Lys58 and the aldehyde group of bombykal (Figure S5). Since all three chemical groups are functionally required for this chemical interaction, it offers multiple avenues to probe this hypothesis.

We first tested the aldehyde requirement by evaluating the binding mode of the partial agonist bombykol, whose terminal alcohol group is not electrophilic (Figure 2C). We solved the cryo-EM structure of BmOR3/BmOrco with bombykol (2.81Å, Figure S2C, S3C, Table S1) and found that, while clear bombykol density occupies the pocket, it is not continuous with either Lys58 or Cys215 (Figure 2B and Figure S3C). Instead, Lys58 establishes a hydrogen bond with the alcohol group of bombykol, and no distinct interaction with Cys215.

Next, we evaluated the receptor’s nucleophiles through site-directed mutagenesis (Figure 2B and Table S2). Mutation of Lys58 to Alanine abolished responses to both bombykal and bombykol. However, substituting Lys58 for Glutamine (which preserves hydrogen bond capabilities but lacks a primary amine) fully restored response to bombykol but not bombykal. This result confirms that the lysine primary amine is strictly required for aldehyde, but not alcohol, recognition. Similarly, mutation of Cys215 to alanine selectively disrupts sensitivity to bombykal, functionally validating its unique role in aldehyde recognition.

It is interesting to note that while the Lys58Ala mutation eliminated bombykal responses entirely, Cys215Ala produced a milder reduction. This result suggests that in the absence of Cys215, Lys58 can still interact with bombykal, but that the converse is not the case. We hypothesize that, in the absence of Cys215, Lys58 may be able to form a Schiff base with the bombykal aldehyde, another type of reversible covalent interaction between aldehydes and Lysines, most famously observed in sensory biology for mediating the interaction between retinal and opsins in the visual system [30, 31]. However, in the absence of Lys58 as a nearby base, Cys215 is less likely to be deprotonated to form the more active thiolate, thereby diminishing its reactivity towards the aldehyde (Figure S5). Ligand recognition by BmOR3 therefore relies on a specific covalent interaction exclusively available to aldehyde ligands.

### A strict geometric requirement for activation of BmOR3 confers selectivity towards the hydrophobic tail of bombykal

Lepidopteran PRs are exquisitely selective not just for the functional headgroup but also for the exact position and stereochemistry of the double bonds in the hydrophobic tail [32]. In our structure, the hydrophobic tail of bombykal tucks neatly into a tunnel oriented towards the S6 helix of BmOR3, a region that has been critically implicated in activation of insect ORs (Figure 2D). In particular, the terminal kink of bombykal’s tail is lined by Leu349 and Val352 in S6 and Phe159 in S3. Alanine substitutions at these three positions significantly attenuate the response to bombykal, indicating that these interactions are critical for stabilizing the open conformation (Figure 2D, Figure S4D, Table S2). In contrast, mutations of neighboring residues Val348 and Leu353 that lack specific contacts with bombykal have no effect on activity (Figure 2D, Table S2).

This strict spatial requirement for engagement of the S6 helix explains why bombykol acts as a partial agonist. Binding of BmOR3 to bombykal through a thio-hemiaminal results in the loss of a water molecule, pulling the bombykal molecule closer to Lys58 and positioning the conjugated diene to optimally engage with Phe159, Leu349 and Val352 (Figure 2E). In contrast, bombykol recognition relies on hydrogen bonding with Lys58, which positions the ligand further away from Lys58. This slight shift in positioning of the functional group of bombykol misaligns the conjugated diene of its hydrophobic tail and fails to optimally engage S3-S6 residues required for activation. The poor fit of the hydrophobic tail of bombykol is therefore a consequence of the binding mechanism at the polar end, and results in suboptimal activation regardless of concentration, explaining its partial agonism.

In sum, the bimodal architecture of the BmOR3 pocket drives pheromone specificity through strict chemical and geometric requirements, reminiscent of the classic ‘lock-and-key’ model proposed over a century ago to explain molecular specificity [33].

### Structural insights enable rational design of small-molecule PR inhibitors

As disease vectors, agricultural pests, or essential pollinators, insect populations are profoundly intertwined with human health and prosperity [10, 13]. Targeted control of insect populations is therefore a global priority. Because broad-spectrum insecticides pose ecological and health risks, modulating insect behavior through pheromone pathways is a highly species-specific strategy that disrupts essential, non-redundant reproductive behaviors, minimizing the rapid evolution of resistance and protecting non-target organisms. While the potential of using pheromones for the control of insect populations has been recognized for decades, its widespread agricultural use remains challenging. Natural pheromones are often expensive to synthesize, chemically unstable, and inherently lack the high volatility desirable for broad field application. Leveraging structural insights into pheromone recognition is a promising inroad towards the rational design of synthetic pheromone analogues with custom-designed properties to act as potent PR inhibitors or activators. This approach would address the pressing need for selective control of insect populations by focusing on behavioral manipulations rather than broad toxicity.

We leveraged our structural insights on BmOR3 to undertake the first rational design of an irreversible small molecule inhibitor of an insect PR. Because our results suggest that proximity to Lys58 in the PR binding pocket likely results in deprotonation of Cys215 into a highly nucleophilic thiolate (Figure 2C), we targeted Cys215 for nucleophilic substitution. We synthesized a series of α-chloroacetamides (ACAs), which are thiolate-specific electrophilic warheads, with saturated aliphatic tails ranging from 4 to 9 carbons (Figure 3A). Using our *in vitro* calcium flux assay in HEK293T cells, we evaluated the dose-dependent inhibition of BmOR3. ACAs proved to be potent inhibitors, with potency scaling with carbon tail length to reach low nanomolar IC_50_s (Figure 3A, Table S3). We speculate that longer tails that better mimic the natural pheromone likely improve potency by anchoring the molecule in the pocket long enough for the warhead to covalently react with Cys215. Importantly, these compounds did not activate BmOR3, because the lack of conjugated dienes in the aliphatic tail of the ACA inhibitors impairs their ability to engage the S6 helix of the PR pocket, thereby acting as irreversible inhibitors.

**Figure 3.**
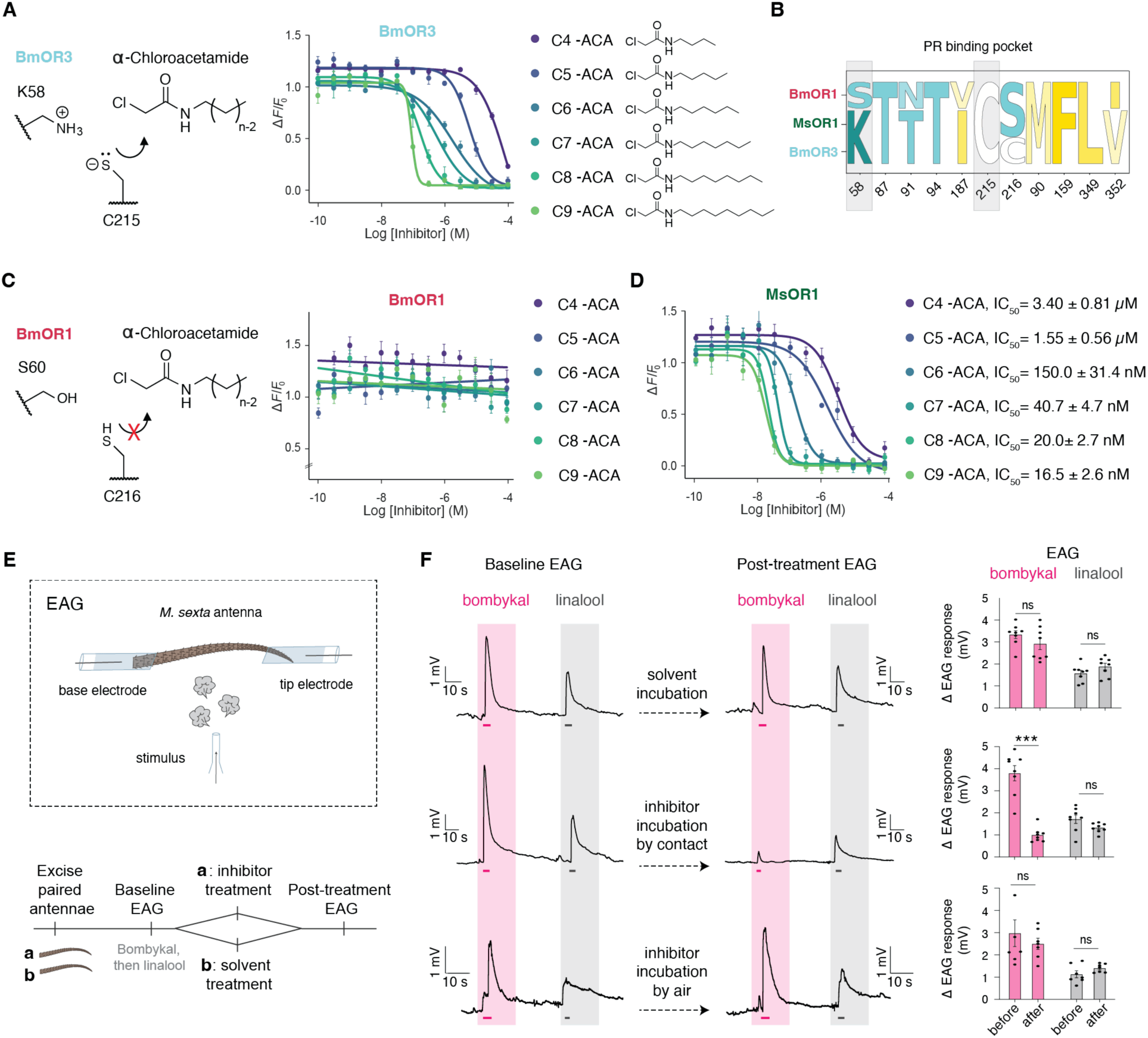
Rationally designed covalent inhibitors selectively inhibit bombykal-sensing PRs *in vitro* and *ex vivo*. (**A**) Left, Schematics of the proposed chemical interaction between alpha-chloroacetamide (ACA) compounds and the Cys215 thiolate in BmOR3, facilitated by the neighboring Lys58 base. Right, Dose-dependent inhibition of BmOR3 by ACAs of varying carbon lengths (activation measured against a fixed bombykal concentration post-incubation). (**B**) Sequence alignment of key binding pocket residues across BmOR1, MsOR1, and BmOR3. Residues are colored by hydrophobicity as in Figure 1 (cyan, polar; gold, hydrophobic). (**C**) Left, Schematic illustrating the lack of ACA reactivity with the Cys 215 in BmOR1 due to the absence of a neighboring lysine to favor thiolate formation. Right, Dose-response curves showing ACAs fail to inhibit BmOR1 (assayed against a fixed bombykol concentration). (**D**) Dose-dependent inhibition of the pest receptor MsOR1 by ACAs (assayed against a fixed bombykal concentration). (**E**) Schematic of the electroantennogram (EAG) recording setup (upper) and the timeline for the paired *ex vivo* treatment of excised *M. sexta* antennae with solvent or ACA (lower). (**F**) Left and middle: Representative EAG traces from *M. sexta* antennae in response to bombykal (magenta) or the control odor linalool (grey) before and after treatment. Dashes indicate 1-s stimulus delivery. Right, Quantification of the change in peak EAG amplitude after treatment for each stimulus. N = 7–8 antennae per group. Bars represent mean ± SEM, dots indicate individual replicates. *** P < 0.001 (two-way repeated measures ANOVA with Sidak’s post hoc test).

To confirm that ACAs specifically target Cys215, we generated alanine mutations for all three cysteines present in the BmOR3 binding pocket and tested them against our most potent compound, ACA-C9 (Figure S6A and Table S3). Only the C215A mutation abolished inhibition, confirming the successful targeting of the intended site of action. To further validate the specificity of ACAs towards an activated cysteine, we tested the ACA series against BmOR1, a related PR from *B. mori* that features a cysteine in the exact orthologous position to Cys215 but lacks the neighboring lysine required to form the reactive thiolate (Figure 3B,C, Table S3). As predicted, the ACA series showed no inhibitory activity against BmOR1. Together, these results demonstrate that structural insights can drive the rational design of potent, highly selective PR inhibitors.

### Translating structural insights into *in vivo* behavioral control

To test whether these structural insights could be leveraged against agricultural pests without further structural determination, we turned to *Manduca sexta* (the tobacco hornworm). The *M. sexta* essential pheromone blend contains a 1:2 ratio of (*E*,*E*,*Z*)- 10,12,14-hexadecatrienal and bombykal, the latter of which is detected by MsOR1 [34–36]. This PR shares ∼60% sequence identity with BmOR3 and, importantly, retains the exact orthologous lysine and cysteine residues in its binding pocket (Figure S6B). We hypothesized that MsOR1 detects bombykal through formation of a thio-hemiaminal and used targeted mutagenesis and *in vitro* calcium flux assays to confirm this hypothesis. Mutation of Lys59 (Lys58 in BmOR3) to Glu and Ala and Cys217 (Cys215 in BmOR3) to Ala all abolished activation of MsOR1 by bombykal (Figure S6B). Having determined the chemical similarity between the pockets of BmOR3 and MsOR1, we predicted that Cys217 in MsOR1 would be highly reactive and shifted towards formation of the thiolate, as shown for Cys215 in BmOR3. We therefore evaluated the effectiveness of our ACA series against MsOR1. As predicted, ACAs proved to also be highly potent inhibitors of MsOR1, reaching nanomolar IC_50_s as carbon tail length increased (Figure 3D and Table S3).

We next tested these inhibitors in an *ex vivo* setting by recording electroantennograms (EAGs, Figure 3E) from antennae excised from adult male *M. sexta* moths. For each antenna, we first determined the baseline response to bombykal (detected by MsOR1) and a control odor linalool (detected by other ORs in the male *M. sexta* antennae that are not MsOR1) (Figure 3E,F). We established robust responses with baselines that were consistently two-fold higher for bombykal than the control odor linalool. To test the ACA inhibitors, we recorded baseline EAG responses from *M. sexta* antennae, and then incubated them by soaking their base in an ACA-inhibitor solution or solvent control. Following inhibitor treatment and rinsing, we returned the antennae to the rig and recorded changes in EAG responses. We found that only the ACA-inhibitor solution and not the solvent control selectively abolished EAG responses to bombykal, with no effect on the response to linalool (Figure 3F).

While direct contact with inhibitor solution yielded potent and selective inhibition of the bombykal response, incubation of antennae with the volatile phase of the ACA-inhibitor solution did not (Figure 3F and Figure S6C). This result suggests that targeting MsOR1 with a potent electrophile that partially mimics the pheromone is a successful chemical strategy for inhibition, but the vapor pressure of ACA is not high enough to penetrate through the volatile phase and reach the PR (Table S4). As the low volatility of ACA is largely contributed by the amide group, we leveraged the modular design of our compound and synthesized an alternative cysteine-targeting inhibitor with a more volatile warhead: a vinyl ketone (Table S4). We tested the effect a 9-carbon vinyl ketone (VK-C9) by conducting *ex vivo* experiments similar to those used for ACA inhibitors, but delivering the inhibitor compound through the volatile phase instead of by soaking. Our results indicate that VK-C9 achieved robust, highly selective inhibition of the bombykal response delivered entirely through the vapor phase (Figure 4A and Fig. S6C).

**Figure 4.**
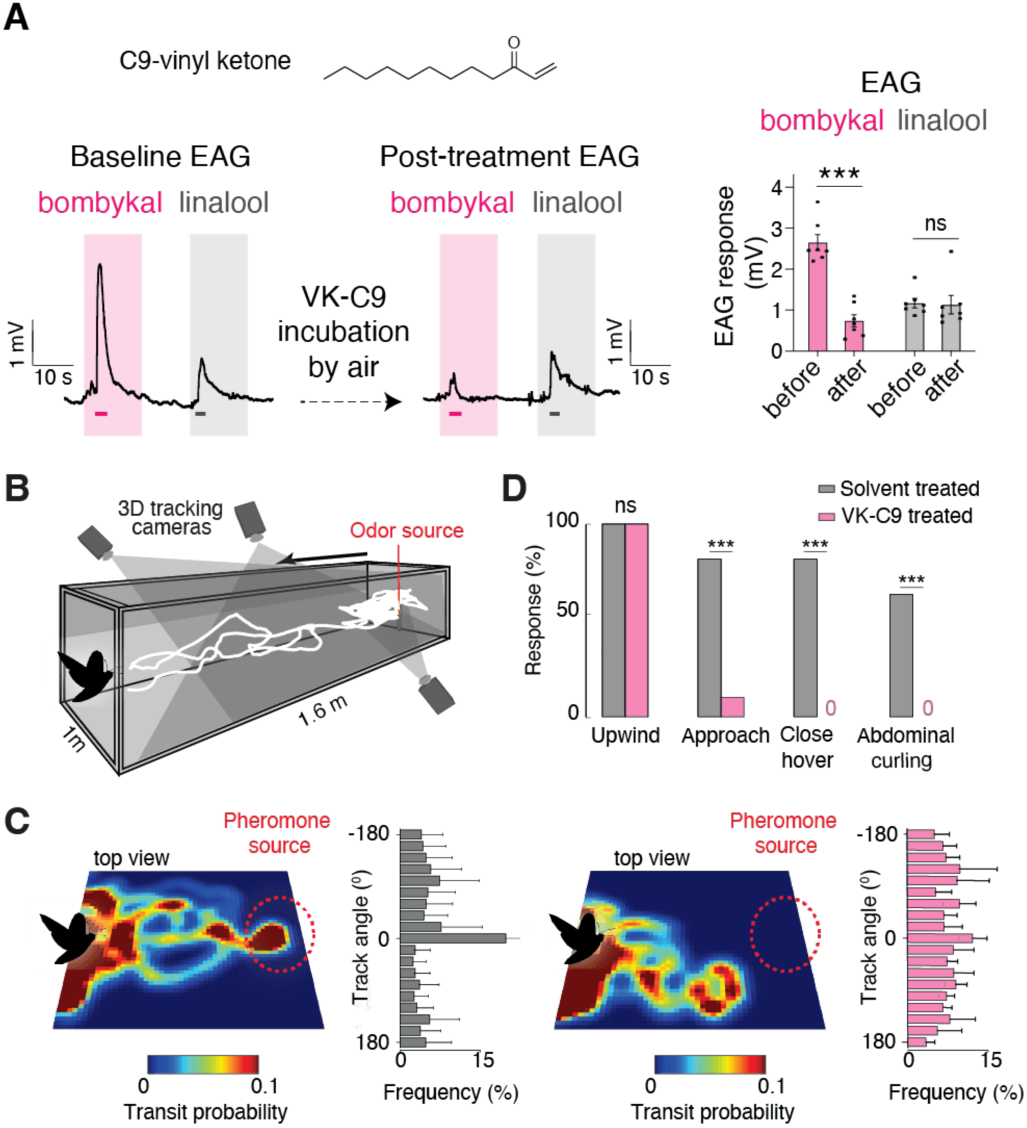
Vinyl ketone inhibitors disrupt pheromone tracking *in vivo*. (**A**) Left and middle: Representative EAG traces from *M. sexta* antennae responding to bombykal (magenta) or the control odor linalool (grey) before (left) and after (middle) exposure to a 9-carbon vinyl ketone (VK-C9). Magenta dashes indicate 1-s stimulus delivery. Right: Quantification of peak EAG voltage changes. n = 7 antennae. Bars represent mean + SEM with individual replicates as dots. *** P < 0.001 (two-way repeated measures ANOVA with Sidak’s post hoc test). (**B**) Schematic of the *M. sexta* wind-tunnel pheromone tracking behavioral assay. (**C**) Representative spatial occupancy heatmaps for male moths pre-exposed to hexane solvent (left) or the volatile phase of VK-C9 (right) and the corresponding quantification of the proportion of flight time directed toward the pheromone source. (**D**) Percentage of solvent treated (gray) versus VK-C9 treated (magenta) moths successfully executing pheromone-tracking behaviors. n = 9-11 moths. The differences among each the treatment for each behavior: upwind flight (χ² = 0, *P* = 1); approach (χ² = 8.144, ** *P* = 0.004); close hover (χ² = 11.172, *** *P* < 0.001); abdomen curling (χ² = 6.897, ** *P* = 0.009).

Having developed a potent, selective and airborne MsOR1 inhibitor, we evaluated its efficacy *in vivo* using a wind tunnel assay to monitor long-range mate-tracking behavior (Figure 4B). We positioned the native pheromone blend at one end of the tunnel and released male moths at the opposite end. Control males displayed robust tracking behavior, characterized by directed flight towards the pheromone source (‘approach’), landing (‘close hover’), and mating attempts (‘abdominal curling’) (Figure 4C,D and Figure S6D). In contrast, brief exposure to the volatile phase of VK-C9, but not to solvent control, completely abolished pheromone tracking behavior without impairing general flight or spatial navigation (Figure 4C,D and Figure S6D).

Ultimately, these results provide a direct bridge from atomic-level structural biology to whole-organism behavioral control. By defining the mechanistic basis of pheromone sensing across a PR cluster, we establish a rationally designed, modular chemical framework that can be tuned to drive highly specific, targeted behavioral interventions.

## Discussion

Fast and unequivocal mate recognition is a critical function of the olfactory system, driven by the exquisite precision of the pheromone sensing pathways. In this work, we elucidate the structural logic of pheromone recognition across the lepidopteran clade, uncovering a unique covalent binding strategy used to detect aldehyde odorants with high selectivity. Leveraging these structural insights, we establish a proof-of-principle pipeline to rationally design small-molecules that exploit the unique chemistry of pheromone receptors to modulate insect behavior. We show that this pipeline can successfully disrupt the sexual communication system of an ecologically relevant insect pest, translating atomic-level receptor mechanisms directly into *in vivo* behavioral control.

The high chemical specificity of BmOR3 represents a sharp divergence from the recognition strategies of the broadly tuned ORs characterized thus far [37, 38]. In promiscuous ORs, ligands rely on flexible, hydrophobic interactions that allow diverse odorants to occupy a variety of poses [37]. Pheromone receptors, in contrast, impose a strict binding mode requiring precise chemical and geometric complementarity, closely resembling the classic lock-and-key paradigm of molecular recognition.

Ligand recognition strategies dependent on covalent bond formation as we observed here are likely broadly employed across the animal kingdom, beyond insect pheromone sensing. Volatile aldehydes are abundant natural odors, and their inherent chemical reactivity offers multiple mechanisms for selective interactions with specific receptors. While most odorant receptors across the tree of life are orphan, many feature buried lysines within their binding pockets [39]. The placement of a basic and reactive amino acid in an unfavorable hydrophobic environment is suggestive of a specific chemical function. Supporting this, a back-engineered mammalian odorant receptor OR6A2 was recently shown to detect its ligands, aliphatic aldehydes, via a reversible Schiff base linkage between the ligands’ aldehyde group and a lysine in the pocket (Lys4.60) [39]. We propose that covalent bond formation between reactive aldehydes and strategically placed nucleophiles in the receptors is a widespread mechanism for volatile aldehyde discrimination across clades.

In current models, olfaction typically relies on the transient binding of odorants to receptors to initiate short-lived neuronal signaling events [38]. While thio-hemiaminals are reversible covalent linkages [40], the establishment of this type of bonds between a pheromone and its receptor might impart a different kinetic signature, potentially extending receptor occupancy and altering the downstream signaling cascade or termination mechanism. This covalent recognition strategy may be a first for the olfactory system, but it has a parallel in the visual system, where a reversible Schiff base formation between the aldehyde group of the chromophore retinal and a buried lysine in the rhodopsin receptor underlie light sensing [30, 31]. Future work connecting the binding kinetics of covalent ligands to neuronal firing rates will illuminate how unique receptor chemistries shape the temporal dynamics of olfactory signaling.

The lock-and-key logic of pheromone sensing we have uncovered suggests a clear strategy for designing custom small-molecule modulators of mating behavior. Historically, modulation of animal behavior through olfaction relied exclusively on empirical trial-and-error, from the discovery of the insect repellent DEET [41] and repellent oils [42] to the use of synthetic pheromone lures [6]. An inability to systematically optimize these tools due to a lack of structural insights has hindered their broad implementation. Our work here now demonstrates that the structural characterization of the insect olfactory system can readily enable the rational design of selective pharmacological modulators, paving the way for next-generation, precision control. Guided by our structural insights, we determined that a highly conserved cysteine in the polar end of lepidopteran binding pockets can be targeted with species-specific precision, as its reactivity is dictated by the unique chemical environment of each specific PR binding pocket. Because of the modularity of the binding pocket, three distinct elements can be independently tuned to dictate specificity and pharmacology. First, the chemical nature of the electrophilic warhead can be modified to achieve desired degrees of reactivity, for example, by substituting halogens in alpha-chloroacetamides or using alternative warheads like acrylamides. Second, the length of the aliphatic tail can be adjusted to maximize binding affinity and inhibitor potency, as demonstrated by the stepwise increases in potency with each added methylene group of our ACA series. Finally, incorporating conjugated dienes at specific locations and with defined stereochemistry could convert an irreversible inhibitor into an irreversible activator, providing a complementary pharmacological strategy for behavior disruption.

Beyond enabling structure-based modulator design, our work demonstrates that engineering of airborne behavior modulators requires a fundamentally distinct physico-chemical profile than that of conventional medicinal chemistry for human therapeutics. Whereas drug-like molecules are typically designed to be water-soluble, crystalline, and three-dimensionally shaped to effectively engage their protein targets in aqueous environments [43], synthetic modulators of olfactory systems must function in the volatile phase. Consequently, they must be substantially less polar, of lower molecular weight, more volatile, and possess a conformationally flexible topology to traverse the air and fit deep within the hydrophobic binding pockets of olfactory receptors. In this work, our modular design framework offers a set of guidelines to accommodate these unique biophysical constraints. Future discovery campaigns aiming to develop new small-molecule behavioral modulators will similarly need to consider these requirements, exploring uncharted chemical space and developing tailored assays that recapitulate the delivery process of the olfactory system.

Because olfaction orchestrates fundamental survival and reproductive behaviors in insects, targeting these pathways with synthetic volatile molecules offers a unique opportunity to manipulate insect behavior to our benefit. Strikingly, nature has already harnessed these tools for behavioral control: vinyl ketones such as the one used in this study, which are highly volatile and reactive electrophiles, are widely found in nature as alarm pheromones and defense compounds to trigger specific behavioral responses [44–46]. Our structure-guided design now makes it possible to build upon these naturally selected strategies to add precision and flexibility for targeting a wide range of receptors, opening the way for the rational design of behavior modulating compounds.

## Acknowledgments

We thank Jaewook Ryu for early contributions to this work and all members of the del Mármol lab for support and feedback. We thank Ashley Su and Melanie Anderson in the Riffell lab for their help with the wind tunnel behavior assay and setting up EAG. We thank Jon Clardy and James Osei-Owusu for stimulating discussions and advice. We thank Marie Bao for feedback on the manuscript. We thank Rui Yan, Nicholas Spellmon, Zhening Zhang at the Janelia CryoEM facility, and Richard Walsh and members of the Structural Biology Center at Harvard Medical School for support with cryoEM data acquisition. We thank the SBGrid Consortium for support. We thank Jennifer Smith and Richard Siu at the ICCB-Longwood Screening Facility for help with GCaMP data collection. J.d.M. is a Freeman Hrabowski Scholar of the Howard Hughes Medical Institute.

## Author contributions

Conceptualization: SSJ, QZ, JdM

Methodology: SSJ, SATM, PKC, MS, SM, XZ, HJ, PAC, JR, QZ, JdM

Investigation: SSJ, SATM, PKC, MS, SM, HJ, XZ, QZ

Visualization: SSJ, SATM, PKC, SM, XZ, JdM

Funding acquisition: JdM

Project administration: SSJ, JdM

Supervision: QZ, JR, PAC, JdM

Writing – original draft: SSJ, JdM

Writing – review & editing: SSJ, SATM, PKC, MS, SM, PAC, QZ, JR, JdM

## Data and materials availability

The cryo-EM density maps and the atomic models of the complexes have been deposited in the Electron Microscopy Data Bank and Protein Data Bank with the following accession numbers EMD-XXXXX and PDB:XXXX (BmOR3/Orco, apo), EMD-XXXXX and PDB:XXXX (BmOR3/Orco, bombykal bound), EMD-XXXXX and PDB:XXXX (BmOR3/Orco, bombykol bound). All other data needed to evaluate the conclusions in the paper are present in the paper and/or the supplementary materials.

