## Supplementary material for "Molecular logic of pheromone recognition enables rational design of insect behavior modulators": SI

### Materials and Methods

Synthesis of Bombykal ((10E,12Z)-hexadeca-10,12-dien-1-al)

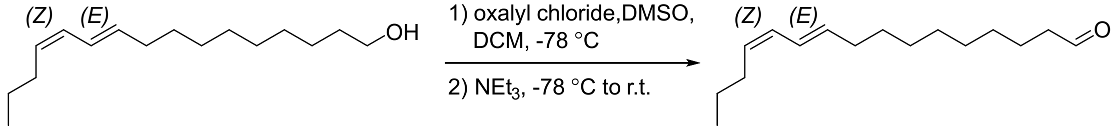


Bombykal was synthesized from commercially available bombykol ((10*E*,12*Z*)-hexadeca-10,12-dien-1-ol, MedChemExpress) following a modified literature procedure [1, 2]. To a solution of oxalyl chloride (8.6 µL, 0.10 mmol, 1.2 eq) in dichloromethane (1.2 mL) was added DMSO (12.0 µL, 0.168 mmol, 2.0 eq) at –78 °C to form a white suspension. After 10 minutes, a solution of bombykol (20 mg, 0.084 mmol, 1 eq) in dichloromethane (0.22 mL) was added dropwise to the reaction mixture at –78 °C and kept stirring for 30 min which resulted in a colorless solution. Triethylamine (70 µL, 0.5 mmol, 6.0 eq) was then added, and the reaction mixture was allowed to warm to room temperature. After 30 minutes, $H_2O$ (10 mL) was added, and the aqueous phase was extracted with dichloromethane (10 mL, repeated three times). The combined organic phase was dried over sodium sulfate, filtered, and concentrated. The residue was purified by flash column chromatography on silica gel (using 1% to 2% ethyl acetate/hexane as eluent) to afford bombykal as colorless oil (99%, 20 mg). $^1$H NMR (500 MHz, $CDCl_3$) δ 9.76 (t, J = 1.9 Hz, 1H), 6.34 – 6.25 (m, 1H), 5.95 (dd, J = 10.9, 10.7 Hz, 1H), 5.65 (dt, J = 14.6, 7.0 Hz, 1H), 5.30 (dt, J = 11.0, 7.6 Hz, 1H), 2.41 (td, J = 7.4, 1.9 Hz, 2H), 2.17 – 2.11 (m, 2H), 2.11 – 2.05 (m, 2H), 1.68 – 1.56 (m, 2H), 1.43 – 1.19 (m, 12H), 0.92 (t, J = 7.4 Hz, 3H). $^{13}$C NMR (126 MHz, $CDCl_3$) δ 203.07, 134.71, 130.04, 128.90, 125.84, 44.05, 32.98, 29.89, 29.50, 29.44, 29.41, 29.29, 29.27, 23.04, 22.21, 13.93. The NMR spectra agree with the reported data [3].

Synthesis of α-chloroacetamide compounds

*Note: C4-ACA, C5-ACA, and C6-ACA were purchased from commercial sources and used without further purification.*

Synthesis of C7-ACA (2-chloro-N-heptylacetamide)

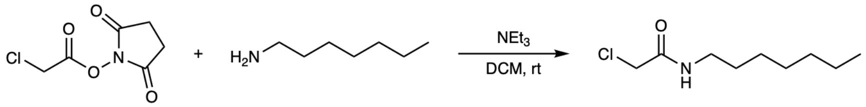


**C7-ACA**

A 1-dram vial equipped with a stir bar was charged with heptan-1-amine (57.6 mg, 0.500 mmol), and 2,5-dioxopyrrolidin-1-yl 2-chloroacetate (114.9 mg, 0.6000 mmol, 1.2 equiv). DCM (5.0 mL) and triethylamine (101.2 mg, 1.000 mmol, 2.0 equiv) were added sequentially. The mixture was stirred at autogenous temperature in ambient environment (20 ºC) for 4 h. The reaction mixture was concentrated *in vacuo*. The residue was dissolved in 0.2 mL of DCM and directly purified by flash chromatography (FlashPure 4-

g Si Cartridge, 0–50% ethyl acetate in hexane linear gradient over 11 min, flow rate 18 mL/min). The title compound was isolated as a white solid (70.1 mg, 0.366 mmol, 73% yield). $^1$H NMR (400 MHz, $CDCl_3$) δ 6.56 (s, 1H), 4.04 (s, 2H), 3.30 (td, $J$ = 7.3, 5.9 Hz, 2H), 1.54 (t, $J$ = 7.2 Hz, 2H), 1.40 – 1.21 (m, 8H), 0.95 – 0.80 (m, 3H). Accurate mass (ESI-TOF) calculated for $C_9H_{19}ClNO$ $(M + H)^+$ 192.1155, found 192.1121.

Synthesis of C8-ACA (2-chloro-N-octylacetamide)

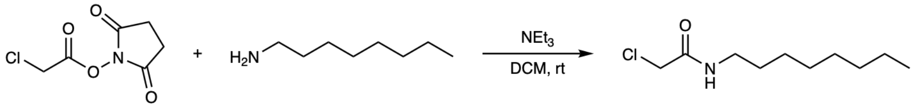


**C8-ACA**

A 1-dram vial equipped with a mini-stir bar was charged with octan-1-amine (65.6 mg, 0.500 mmol), and 2,5-dioxopyrrolidin-1-yl 2-chloroacetate (114.9 mg, 0.6000 mmol, 1.2 equiv). DCM (5.0 mL) and triethylamine (101.2 mg, 1.000 mmol, 2.0 equiv) were added sequentially. The mixture was stirred at autogenous temperature in ambient environment (20 ºC) for 4 h. The reaction mixture was concentrated *in vacuo*. The residue was dissolved in 0.2 mL of DCM and directly purified by flash chromatography (FlashPure 4-g Si Cartridge, 0–50% ethyl acetate in hexane linear gradient over 11 min, flow rate 18 mL/min). The title compound was isolated as a white solid (68.7 mg, 0.334 mmol, 67% yield). $^1$H NMR (400 MHz, $CDCl_3$) δ 6.56 (s, 1H), 4.04 (s, 2H), 3.30 (td, $J$ = 7.3, 5.9 Hz, 2H), 1.53 (q, $J$ = 7.1 Hz, 2H), 1.41 – 1.18 (m, 10H), 0.94 – 0.80 (m, 3H). Accurate mass (ESI-TOF) calculated for $C_{10}H_{21}ClNO$ $(M + H)^+$ 206.1311, found 206.1376.

Synthesis of C9-ACA (2-chloro-N-nonylacetamide)

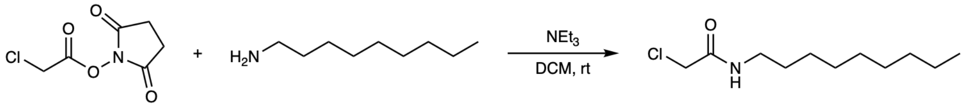


**C9-ACA**

A 1-dram vial equipped with a mini-stir bar was charged with nonan-1-amine (71.6 mg, 0.500 mmol), and 2,5-dioxopyrrolidin-1-yl 2-chloroacetate (114.9 mg, 0.6000 mmol, 1.2 equiv). DCM (5.0 mL) and triethylamine (101.2 mg, 1.000 mmol, 2.0 equiv) were added sequentially. The mixture was stirred at autogenous temperature in ambient environment (20 ºC) for 4 h. The reaction mixture was concentrated *in vacuo*. The residue was dissolved in 0.2 mL of DCM and directly purified by flash chromatography (FlashPure 4-g Si Cartridge, 0–50% ethyl acetate in hexane linear gradient over 11 min, flow rate 18 mL/min). The title compound was isolated as a white solid (100.5 mg, 0.4573 mmol, 91% yield). $^1$H NMR (400 MHz, $CDCl_3$) δ 6.56 (s, 1H), 4.04 (s, 2H), 3.30 (td, $J$ = 7.3, 5.9 Hz, 2H), 1.60 – 1.47 (m, 2H), 1.29 (dt, $J$ = 15.8, 4.1 Hz, 12H), 0.97 – 0.80 (m, 3H). Accurate mass (ESI-TOF) calculated for $C_{11}H_{23}ClNO$ $(M + H)^+$ 220.1468, found 220.1546.

Expression and purification of BmOR3/BmOrco complex

The coding sequence of BmOrco was a gift from Leslie Vosshall (Addgene plasmid #73926; http://n2t.net/addgene:73926 ; RRID:Addgene_73926). The coding sequence of BmOR3 was codon optimized and synthesized from Twist Bioscience. Gene fragments of full-length BmOR3 was cloned into a pEG BacMam vector containing an N-terminal superfolder GFP and an HRV 3C protease site. The gene fragment of full length BmOrco was cloned into a pEG BacMam vector containing an N-terminal mCherry and an HRV 3C protease site. Plasmids were transfected (1:2 ratio of BmOR3 to BmOrco, 750 μg per L) into Expi293F GnTI− cells grown in Expi293 medium with FectoPro (800 μL per L of culture) and kept at 37 °C with 8% carbon dioxide. After 18-24 h ,3 mM valproic acid and 0.5 % glucose were added, and the temperature was dropped from 37 °C to 30 °C for an additional 48-72 hours. Cells were then pelleted by centrifugation, flash frozen, and stored at −80 °C until the day of purification.

For purification, cell pellets were resuspended in 100 mL of ice cold solubilization buffer per liter of cell culture. The solubilization buffer was composed of 20 mM HEPES/NaOH (pH 7.5), 150 mM NaCl, 0.5% (w/v) Lauryl Maltose Neopentyl Glycol (LMNG; Anatrace), 0.1% (w/v) cholesterol hemisuccinate (CHS; Sigma-Aldrich), 1 μg/mL leupeptin, 1mM benzamidine, 1 μg/mL aprotinin, and 1 μg/mL pepstatin A, and 1 mM phenylmethylsulfonyl fluoride (PMSF). The cells were kept in the cold room while rotating for 2 hours to extract the membrane protein complexes from the cell membranes to the micelle environment. The mixture was then clarified by centrifugation at 175,000 x g for 40 minutes, and the supernatant was added to 1 mL of anti-GFP nanobody-coupled Sepharose resin (bead volume) [4] per liter of cell culture. After washes with the Equilibration buffer (20 mM HEPES pH 7.5, 150 mM NaCl, 0.002% LMNG, 0.0004% CHS), the OR/Orco complex was eluted by mixing 50 μg of 3C protease (Sigma-Aldrich) with every 1 mL of resin and gentle rotation at 4 °C for an hour. The sample was then concentrated and injected into a Superose 6 Increase column (Cytiva) pre-equilibrated with the Equilibration buffer. Peak fractions containing BmOR3/Orco complex were pooled and concentrated to A280 = 2.74. The proteins were flash-frozen and stored at −80 °C until use.

CryoEM sample preparation and data acquisition

To prepare the bombykal-bound sample, bombykal was added to the OR3/Orco sample to a final concentration of 20 μM and incubated for an hour at room temperature. The apo sample omitted this step and was directly used for freezing grids. Cryo-EM grids were frozen using a Vitrobot Mark IV (FEI) using the following procedure: 4 μL of the sample was applied to a glow-discharged UltraAufoil R1.2/1.3, 300 mesh gold grid, blotted for 5 s in 100% humidity at 4 °C and Blot force: 1, and plunge frozen in liquid ethane cooled by liquid nitrogen.

Cryo-EM data were recorded on a 300-kV Titan Krios G3i microscope (ThermoFisher), equipped with a Gatan BioQuantum GIF/ K3 direct electron detection camera at the Janelia Research Campus Cryo-EM Facility. SerialEM was used for automated data collection. The defocus range is -1.0 to -2.0 µm. Movies were collected at a magnification of 81,000x in superresolution mode with a physical pixel size of 1.061 Å/pixel. All data was collected with a 30° tilted stage. 50 frames were collected with a total dose of 50 electrons per $Å^2$. Further details of the data collection parameters are listed in Supplementary Table 1.

<u>CryoEM data processing</u>

*BmOR3/Orco with Bombykal*: A total of 6,320 movies were collected for the OR3 sample with bombykal. The movies were aligned using motion corrected algorithms in CryoSPARC [5], and Blob-based autopicking in CryoSPARC [5] was used to select initial particles with 2-fold binning, resulting in 4,498,756 particles. From the initial 2D classification, a total of 2,934,701 particles were selected to reconstruct the initial map of the BmOR3/Orco complex by running an Ab-initio job followed by a homogeneous refinement job. Using this reconstruction as a starting point, a seed-based method previously described in [6, 7], which involves iterative rounds of Ab-initio reconstruction and heterogeneous refinement were ran to remove false-positive particles and rescue 'good' particles that were excluded during the initial 2D classification. This resulted in a stack of 3,283,431 particles. Following non-uniform refinement, the particles were then classified through 3D classification in CryoSPARC [5] using a focused mask on the OR subunit. The 3D classification classified the particles to subsets representing two open-pore classes and two closed-pore class. Further processing of one of the open-pore classes (1,070,550 particles) through non-uniform refinement and reference-based motion correction generated a map with an overall resolution of 2.68 Å with clear bombykal density.

*BmOR3/Orco with Bombykol*: A total of 9,768 movies were collected for the OR3 sample with bombykal. The movies were aligned using motion corrected algorithms in CryoSPARC [5], and Blob-based autopicking in CryoSPARC [5] was used to select initial particles with 2-fold binning, resulting in 6,058,000 particles. From the initial 2D classification, a total of 3,645,202 particles were selected to reconstruct the initial map of the BmOR3/Orco complex by running an Ab-initio job followed by a homogeneous refinement job. Using this reconstruction as a starting point, a seed-based method previously described in [6, 7], which involves iterative rounds of Ab-initio reconstruction and heterogeneous refinement were ran to remove false-positive particles and rescue 'good' particles that were excluded during the initial 2D classification. This resulted in a stack of 3,280,138 particles. Following non-uniform refinement, the particles were then classified through 3D classification in CryoSPARC [5] using a focused mask on the OR subunit. The 3D classification classified the particles to subsets representing two open-pore classes and two closed-pore class. Further processing of one of the open-pore classes (1,191,576 particles) through non-uniform refinement and reference-based motion correction generated a map with an overall resolution of 2.76 Å with clear bombykol density.

*BmOR3/Orco without ligand:* A total of 5,389 movies were collected for the OR3 sample in the absence of ligand. The movies were aligned using motion corrected algorithms in CryoSPARC [5], and Blob-based autopicking in CryoSPARC [5] was used to select initial particles with 2-fold binning, resulting in 4,652,403 particles. From the initial 2D classification, a total of 2,543,759 particles were selected to reconstruct the initial map of the BmOR3/Orco complex by running an Ab-initio job followed by a homogeneous refinement job. Using this reconstruction as a starting point, a seed-based method previously described in [6, 7], which involves iterative rounds of Ab-initio reconstruction and heterogeneous refinement were ran to remove false-positive particles and rescue 'good' particles that were excluded during the initial 2D classification. This resulted in a stack of 2,445,450 particles. Following non-uniform refinement, the particles were then classified through 3D classification in CryoSPARC [5] using a focused mask on the OR subunit. The 3D classification classified the particles to subsets representing two open-pore classes and two closed-pore class. Further processing of one of the closed-pore classes (327,260) that includes an additional round 3D classification, non-uniform refinement and local refinement generated a map with an overall resolution of 2.89 Å.

Model Building

The Alpha Fold models [8, 9] of BmOrco (AF-Q7YT34-F1-v4) and BmOR3 (AF-Q5FBE0-F1-v4) were used as a starting point for manual model building in Coot [10]. Individual amino acid residues were assigned based on the quality of the side-chain densities in the primary maps.

The structure of bombykal bound, open conformation of BmOR3/Orco was built using both the sharpened and unsharpened maps. For the BmOR3 subunit, N-terminal residues 1-3 and the loop residues connecting S4-S5, 254-269 were omitted due to poor density in the map. For the Orco subunits, N-terminal residues 1-6 and intracellular S4-S5 loop residues 248-317 were omitted due to poor density in the map. Atomic coordinates were refined against the sharpened map using real space refinement implemented in PHENIX [11] for 5 macrocycles with secondary structure restraints applied and without symmetry enforced. The bombykal bound model was refined including the ligand, with restraints obtained using eLBOW implemented in Phenix [11]. A separate edits parameter file was supplemented during the refinement to manually define the covalent linkage between bombykal and Lys58 and Cys215.

The structure of bombykol bound, open conformation of BmOR3/Orco was built using both the sharpened and unsharpened maps. For the BmOR3 subunit, N-terminal residues

1-3 and the loop residues connecting S4-S5, 255-269 were omitted due to poor density in the map. For the Orco subunits, N-terminal residues 1-6 and intracellular S4-S5 loop residues 248-317 were omitted due to poor density in the map. Atomic coordinates were refined against the sharpened map using real space refinement implemented in PHENIX [11] for 5 macrocycles with secondary structure restraints applied and without symmetry enforced. The bombykal bound model was refined including the ligand, with restraints obtained using eLBOW implemented in Phenix [11].

The structure of the apo, closed conformation of BmOR3/Orco was built using both the sharpened and unsharpened maps. For the BmOR3 subunit, N-terminal residues 1-4 and the loop residues connecting S4-S5, 255-268, and the loop residues connecting S3-S6, 338-339 were omitted due to poor density in the map. For the Orco subunits, N-terminal residues 1-6, and intracellular S4-S5 loop residues 248-317 were omitted due to poor density in the map. Atomic coordinates were refined against the density-modified map using real space refinement implemented in PHENIX [11] for 5 macrocycles with secondary structure restraints applied and without symmetry enforced.

All structural biology software were compiled by SBGrid [12].

Pore analysis

Pore diameter along the central axis and side exits were calculated using HOLE [13]. For each structure, separate calculations were conducted for the central pore and the lateral conduits by varying the starting position and the vector defining the orientation of the pore finding.

GCaMP6 fluorescence calcium flux assay

The assay was performed similarly to method descriptions in previous studies [7, 14, 15]. All DNA constructs used in this assay were cloned into a modified pME18 vector that contains a SV40 promotor.

Each transfection condition (per well) contained a total of 300 ng of plasmids in 1:1:1 ratio of the GCaMP6, Orco, and the corresponding OR plasmid that was diluted in 4.8 uL of OptiMEM (Gibco) along with 0.1 uL of P3000. The mixture was incubated for 5 minutes and then further incubated for 20 minutes after being mixed with Lipofectamine 3000 (Invitrogen) diluted in OptiMEM. This mixture was scaled accordingly to batch transfect 50-100 wells per construct per plate.

HEK293T cells were maintained at 37°C with 5% carbon dioxide and grown in high glucose DMEM enhanced with 10% FBS and 1% GlutaMAX (Gibco). Cells were detached with TrypLE Express Enzyme and resuspended in FluoroBrite DMEM (Gibco) with 10% FBS and 1% GlutaMAX (Gibco) to a concentration of $5 \times 10^5$ cells/ml. Cells were combined with each transfection condition and added to 2x16 wells of a 384-well plate

(Grenier CELLSTAR). Cells were kept at 37°C for 18-20 hours before being used for fluorescence plate reading.

Odorant plates were prepared using D300e (Hewlett Packard), a digital, non-contact dispenser. Briefly, stock solutions of bombykol, bombykal, and covalent inhibitors were prepared in DMSO and plated onto 384-plates (Greiner, Catalog #784201) in a titrating volume to obtain a concentration series ranging between 63.2 nM and 20 mM with a final volume of 0.4 μL. The odorant plates were kept at −20 °C until usage.

On the day of the GCaMP6 assay, Bravo (Agilent), a versatile liquid handler was used to prepare both the odorant plate and the assay plate (transfected cell plate) prior to the fluorescence readout using the FDSS7000EX kinetic plate Imager (Hamamatsu). For the odorant plate, the DMSO dissolved stock solutions were diluted in 39.6 μL of Reading buffer (20 mM HEPES/NaOH (pH 7.4), 1× HBSS (Gibco), 3 mM $Na_2CO_3$, 1 mM $MgSO_4$, and 5 mM $CaCl_2$) to give a final concentration ranging from 0.632 nM to 0.2 mM with 1% DMSO. For the cell plate, the FluoroBrite DMEM (Gibco) was exchanged out with the Reading buffer.

For inhibition experiments, inhibitor plates with the DMSO dissolved stock solutions were diluted in 39.6 μL of FluoroBrite DMEM (Gibco) to give a final concentration ranging from 316 nM to 100 mM. Using Bravo (Agilent) 4.4 μL of the inhibitor solutions were added to the cell plate and incubated for 2 hours prior to fluorescence readout. Prior to the experiment, the FluoroBrite DMEM (Gibco) was exchanged out with the Reading buffer.

The kinetics of the fluorescence change upon odorant addition were measured using the FDSS7000EX kinetic plate Imager (Hamamatsu). Excitation was at 480 nm and emission was recorded at 540 nm. The exposure time was set to 0.5 s and the LED power was set to 400 mA. After 20 s of baseline recording, 20 μL of odorant solution was added to each well of the assay plate containing 20 μL of the Reading buffer followed by brief mixing, and the fluorescence of each well was continuously recorded for additional 5-10 minutes. All recordings were carried out at room temperature.

Each concentration of ligand was applied to two technical replicates, which were averaged and considered a single biological replicate. The baseline fluorescence, $F_0$, was determined using the first 10 s of GCaMP6 recording, prior to the addition of ligands. $\Delta F$ was calculated by taking the average of F values in the last 10 s of the fluorescence time trajectory and subtracting by the $F_0$. $\Delta F/F_0$ was then calculated for each well and further corrected by subtracting the $\Delta F/F_0$ obtained from three control wells of the same transfection condition but without any ligand addition. These normalized $\Delta F/F_0$ were averaged across the biological replicates to obtain the representative $\Delta F/F_0$ for each concentration point of the specific construct. Finally, custom written python scripts were used to fit the dose-response curves to the four-parameter Hill equation, from which $EC_{50}$,

$IC_{50}$ and hill coefficient values were extracted. Max ΔF/F was defined as the ΔF/F at the highest concentration of the respective ligand in each dose response curve.

Sequence conservation across Lepidopteran ORs

From the uniprot database, FASTA files for Lepidopteran ORs were downloaded with the following search parameters: Taxonomy 7088, OR lengths between 350-500, odorant receptor, olfactory receptor, NOT coreceptor. This list was then combined with a list of uniprot accession codes with candidate PRs that were identified in [16]. Duplicate OR fasta files were removed using CD-HIT [17] by removing overlapping sequences with sequence similarity over 90%. The filtered set of uniprot IDs were then searched in the alphafold database to obtain 1366 unique Lepidopteran OR PDB files.

We then used FoldMason [18] to generate a multiple structural alignment (MSTA) of these receptors. Phylogenetic tree was then built using IQ-Tree2 with 1000 ultrafast bootstrap replicates [19]. The tree was visualized with iTOL(Interactive Tree of Life) [20]. A single cluster, of 249 ORs which contained 180 of the known PRs was annotated as the PR cluster. These receptors were then subsequently re-aligned in FOLDMASON [21] to output the MSTA file used for observing sequence conservation pattern across the binding pocket residues in Lepidopteran PRs. The average Local Distance Difference Test (LDDT) score, a metric of the quality of the structural alignment, for all core domains of these PRs was 0.849, meaning that they share a highly conserved fold and conservation across residues within these domains can be assessed confidently.

Insect rearing

*Manduca sexta* (tobacco hornworms) was raised from larvae (Great Lakes Hornworm, USA) to adults under standard laboratory conditions [22]. They were kept on a 16 h/8 h (light/dark) cycle at 26°C and given a wheatgerm-based diet (Great Lakes Hornworm, USA). Only male moths were selected to eclose and given access to 10% sucrose. For EAG experiments adult male moths aged 1-5 days were used.

Electroantennography and odor delivery

The electroantennography protocol was adapted from a previous study [23]. An antennae that was cut at the base and tip was placed into an electrode (silver wire in a capillary tube filled with 0.1 M KCl) on each side. Voltage recordings were made using an amplifier (Axopatch 200b, Axon Instruments) and digitizer (Digidata 1550b, Axon Instruments).

Olfactory stimulus was prepared by pipetting 3 ul of 100 mM bombykal or neat linalool (Sigma-Aldrich) onto a piece of Whatmann No. 1 filter paper that was inserted into a glass pipette. The glass pipette was positioned close to the tip of the antennae and attached to the odor delivery system. A puff of carbon filtered air was delivered through the glass pipette for 1 s at a rate of 145 ml/per min under the control of a solenoid valve (The Lee

Company). A vacuum was placed behind the preparation to remove any residual odors after the puff. Pilot recordings showed that air alone puffs did not cause responses.

For by contact treatments cut antennae (tip and base) were dipped into either 10 ul solvent or inhibitor and left with the base exposed for 30-40 mins in an eppendorf tube. For air treatments, 20 ul of solvent or 100 mM inhibitor stocks were pipetted onto filter papers and a constant air stream was passed through the filter paper at 145 ml/min for 10 minutes.

pCLAMP 11 software was used for data acquisition and analysis. All EAG traces were baseline corrected and peak voltage changes acquired. Prism 11 was using for graphing and statistics. A two-way repeated measures ANOVA for each treatment was run before proceeding with Sidak's post hoc test. *$p<0.05$; ** $p<0.01$ and *** $p<0.001$.

<u>Manduca sexta pheromone tracking wind tunnel assay</u>

*Wind tunnel and stimuli*: A Plexiglass wind tunnel (L x W x H = 2 x 1 x 1 m) was used to examine the male moth (n = 30) upwind flight behavior in response to pheromone gland extract or solvent control (Figure 5b). The airflow in experiments was maintained at a longitudinal wind speed of 50 cm/s using a fan that directed the air through the filters and honeycomb baffles before entering the wind tunnel working section. At the beginning of scotophase, naive adult male moths (2- 3 days post eclosion) were individually placed inside the cylindrical wire mesh release cage (L= 28 cm, ∅ = 14.5 cm) positioned 30 cm above the tunnel floor at the downwind end. The moth was released approximately 2 m from the odor source. The odor source was positioned 16 cm downstream of the upwind honeycomb and 40 cm above the windtunnel floor.

The experiment was conducted for 3-4 hours of scotophase. The odor source consisted of a moth-shaped paper model (6.5 cm-by-4 cm) and a Whatman filter paper attached to the abdominal regions, onto which the odor stimulus was applied. Each moth was allowed to fly freely inside the wind tunnel for 2 min while its behavior was recorded. Experiments were conducted under the dim white illumination of (0.75 lux) provided from above the tunnel. Infrared backlighting was used to enable video recording of flight trajectories and behavioral observations. Moths that failed to fly within 2 min of release were excluded from the analysis. Two types of behavioral data were collected during experiments: (1) three-dimensional video acquisition for motion analysis of flight trajectories for each treatment group, and (2) manual scoring of behavioral responses. Behavioral responses were categorized as: (i) upwind flight (moth flying to within 0.5 m of odor source), (ii) approach (moth directed toward the source), (iii) close hover (moth hovering within 10 cm of odor source), and (iv) abdomen curl (pre-copula mating posture). The categorical behavioral responses were compared between the treatments using 2-by-2 $\chi 2$-tests. Statistical significance was determined using an $\alpha$-level of 0.05.

*Video acquisition:* The flight trajectories were recorded at 60 frames per sec using two synchronised Blackfly S USB3 cameras (1.3 MP, Mono, C-mount, Figure 5b) equipped with Tamron M118FM08 lens. An overhead camera was positioned 2.24 m above the wind tunnel floor, providing a field of view of 1.64 m, whereas a second camera was mounted on the downwind side of the tunnel at a height of 1.86 m. Behavioral scoring was assisted by an additional monochrome Point gray camera (Mono, FMVU-O3MTM-CS) equipped with Tokina Lens (6.6 mm, F 1.6) positioned outside the upwind end of the tunnel.

*Pheromone Extraction:* Female sex pheromone extract was prepared from calling 1-4-day-old virgin females as described [24]. Briefly, pheromone glands from 10 females were excised and sequentially extracted in hexane by immersion in 100 µl hexane followed by rinsing in an additional 100 µl hexane. The extract was concentrated and reconstituted to a final volume of 250 µl, corresponding to a concentration of 2 female gland equivalents per 250 µl. The solution is stored at -80$^o$ C in an amber colored bottle until use. For the behavioral experiments, 50 µl of the extract was applied to a filter paper for testing.

*Experimental setup*: The moth's antenna was stimulated with the vinyl ketones-C9 inhibitor (n = 10) or the solvent (hexane) control (n = 11) inside the plastic cups. For the vinyl ketones-C9 inhibitor pre-exposure, 30 µl of 100mM solution was applied to the filter paper inside the pasteur pipette with the continuous airstream directed at each antenna for 20 min at a flow rate of 150 ml/min. To assess moths' non-pheromonal behaviors, another control was setup, whereby moths were pre-exposed to hexane as described above, but their behavior towards hexane was assessed instead of the pheromone (n = 9).

### Figure S1

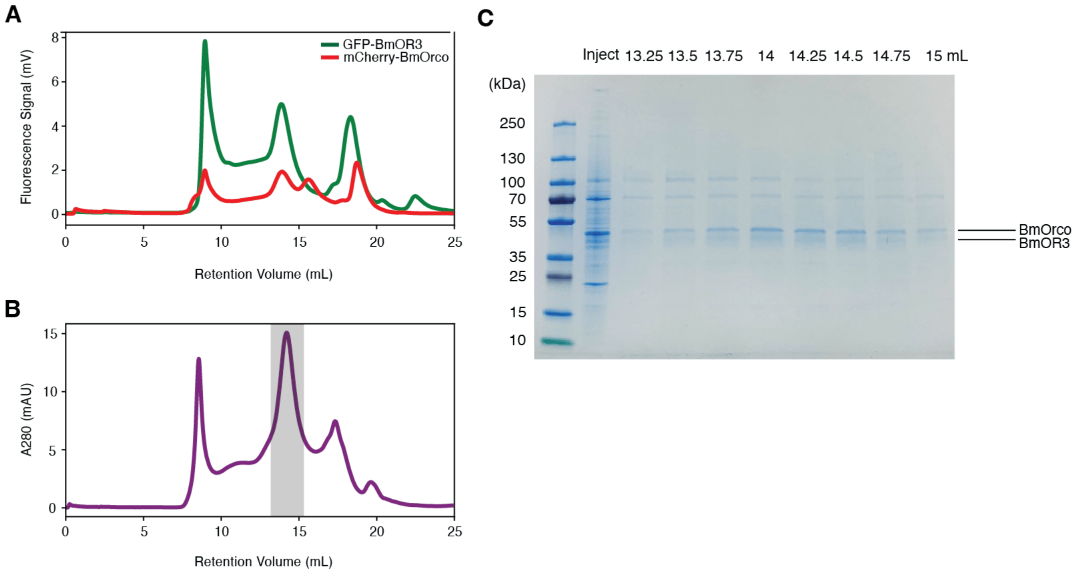


**Figure S1. Purification of BmOR3/BmOrco complex**. (**A)** Fluorescence size-exclusion chromatography (FSEC) profiles of solubilized Expi293F GnTI− cells transiently co-expressing GFP-BmOR3 and mCherry-BmOrco. **(B)** Preparative size-exclusion chromatography (SEC) profile of the BmOR3/BmOrco purified from Expi293F GnTI− cells. Fractions pooled for cryo-EM are highlighted in grey. **(C)** Sodium dodecyl sulfate-polyacrylamide gel electrophoresis (SDS-PAGE) of SEC fractions. First lane contains the molecular weight ladder. Lane labeled as 'Inject' contains a fraction of the sample injected into the SEC column. Remaining lanes correspond to fractions collected at labelled retention volumes. Bands corresponding to BmOR3 and BmOrco are labeled.

### Figure S2

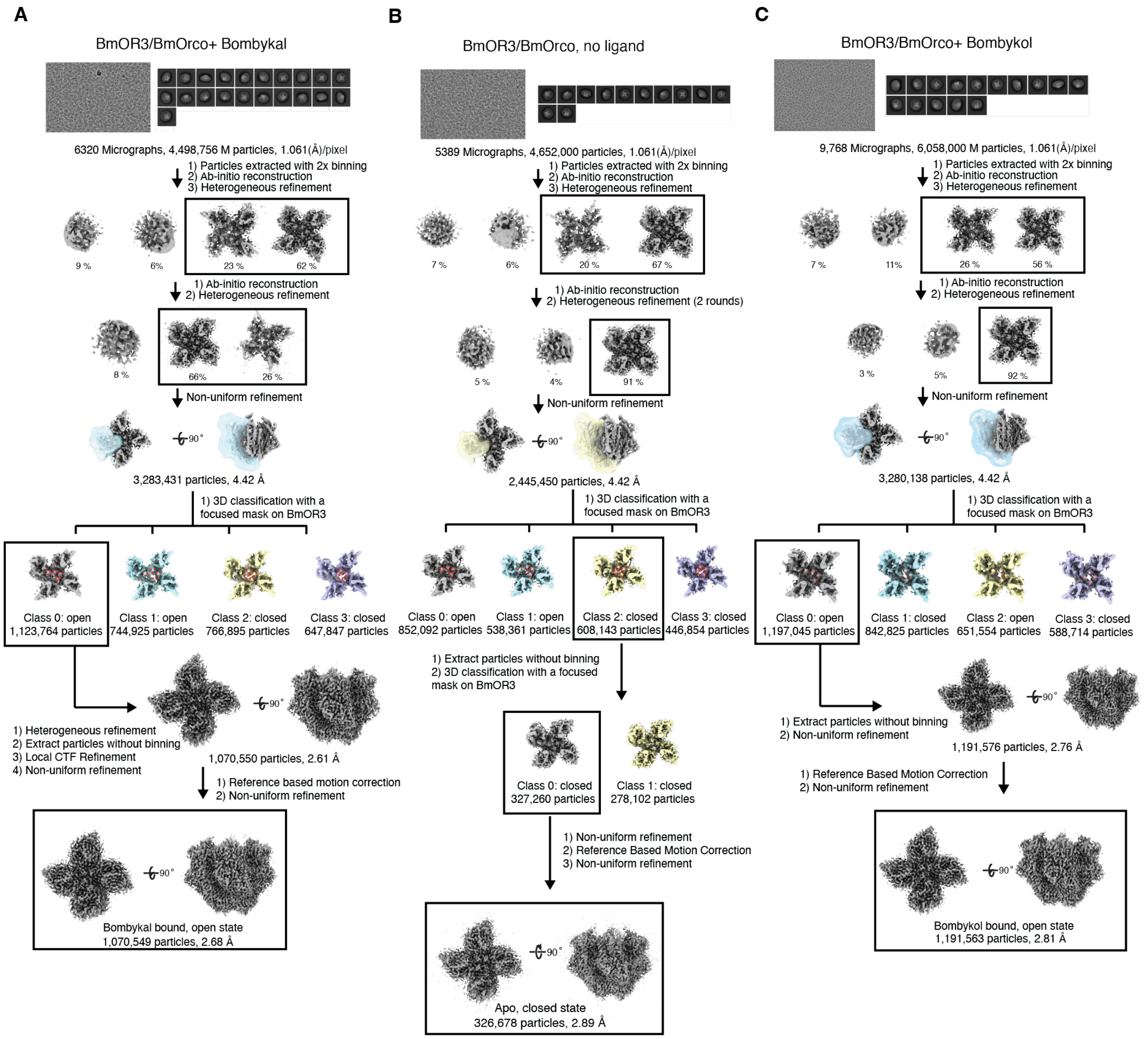


**Figure S2.** Cryo-EM data processing pipelines of BmOR3/BmOrco datasets **(A)** in the presence of bombykal, **(B)** in the apo (unliganded) state, and **(C)** in the presence of bombykol.

### Figure S3

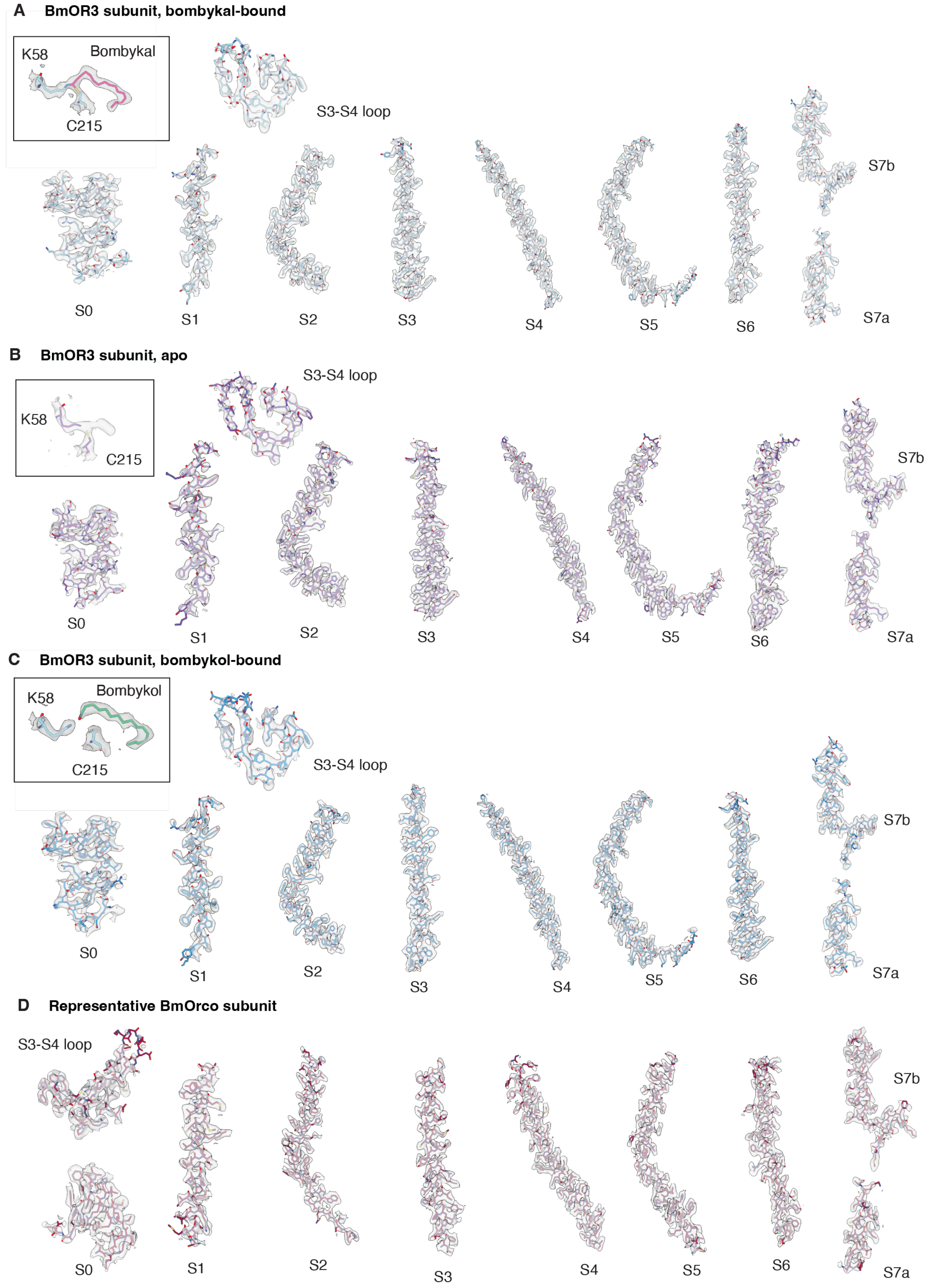
A BmOR3 subunit, bombykal-bound
K58
Bombykal
C215
S3-S4 loop
S0
S1
S2
S3
S4
S5
S6
S7b
S7a
B BmOR3 subunit, apo
K58
C215
S3-S4 loop
S0
S1
S2
S3
S4
S5
S6
S7b
S7a
C BmOR3 subunit, bombykol-bound
K58
Bombykol
C215
S3-S4 loop
S0
S1
S2
S3
S4
S5
S6
S7b
S7a
D Representative BmOrco subunit
S3-S4 loop
S0
S1
S2
S3
S4
S5
S6
S7b
S7a

**Figure S3. Local cryo-EM densities of BmOR3/BmOrco structures.** Cryo-EM density for individual segments of the BmOR3 subunit in the **(A)** bombykal-bound, **(B)** apo, and **(C)** bombykol-bound states. **(D)** Representative cryo-EM density of a BmOrco subunit

### Figure S4

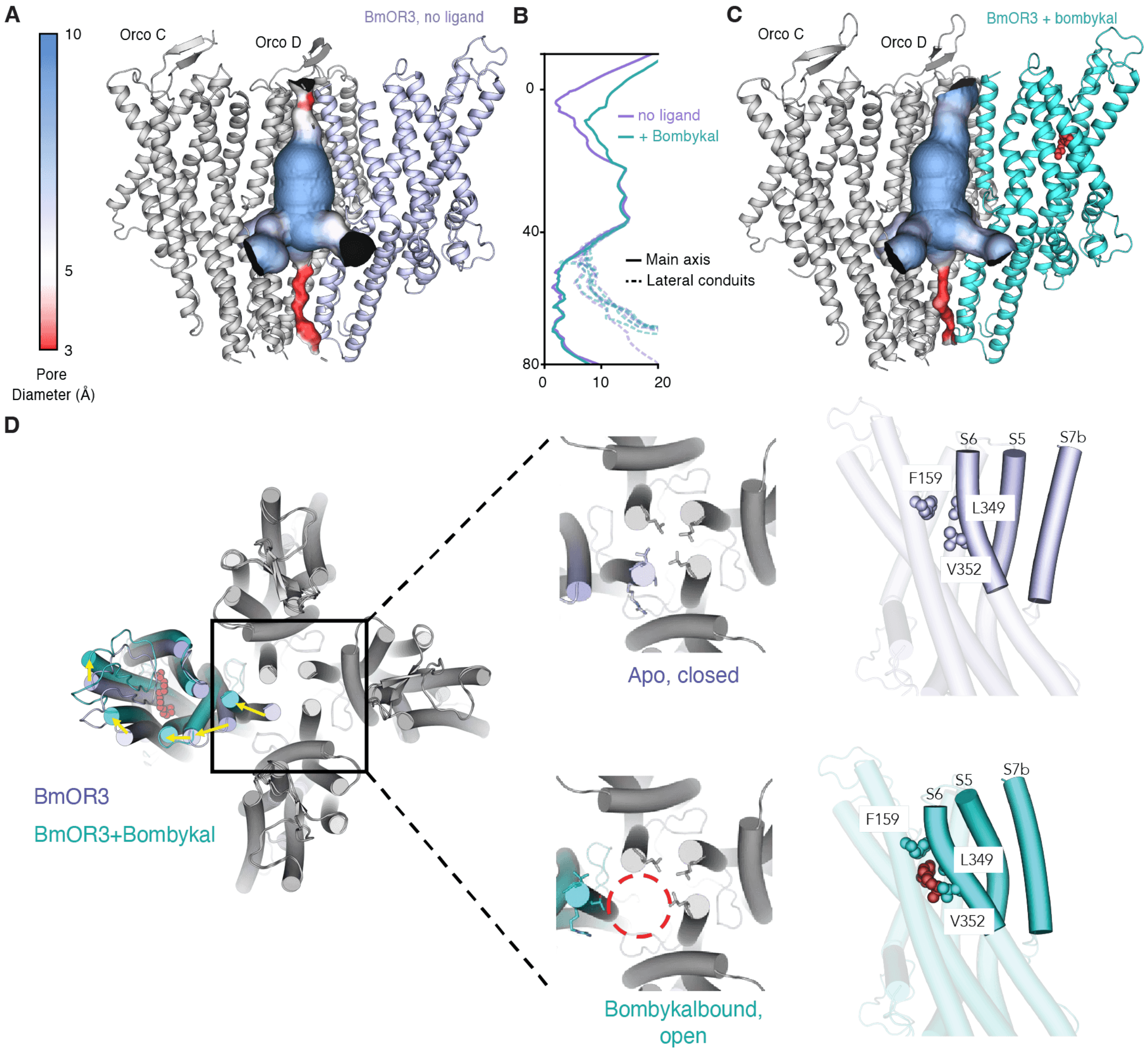


**Figure S4. Structural details of the BmOR3/BmOrco pore.** The ion permeation pathways of the **(A)** apo and **(C)** bombykal bound structures, colored by the pore diameter. The front Orco B subunit is not shown to permit visualization of the central cavity. **(B)** Plot detailing the diameter of the ion permeation pathway along the central axis, measured from the edge of the extracellular side to the intracellular side. BmO3/BmOrco exhibits the quadrivial pore architecture characteristic of this family of receptors. The diameter of the impermeable central pathway through the anchor domain is shown in solid line, whereas those of the four lateral conduits are shown in dashed lines. **(D)** Overlay of top views of the apo and bombykal-bound BmOR3/BmOrco complex, displaying the structural rearrangements associated with receptor activation. Middle inset, focused comparison of structural differences in the pore region of BmOR3 between the resting (top) and activated state (bottom). Right inset, a close-up lateral view of S6, S5, S7b helices in BmOR3 in its resting (top) and activated (bottom) states. Key residues (Leu349 and V352 in S6, F159 in S3) that interact with the hydrophobic tail of bombykal and mediate activation are highlighted.

**Figure S5**

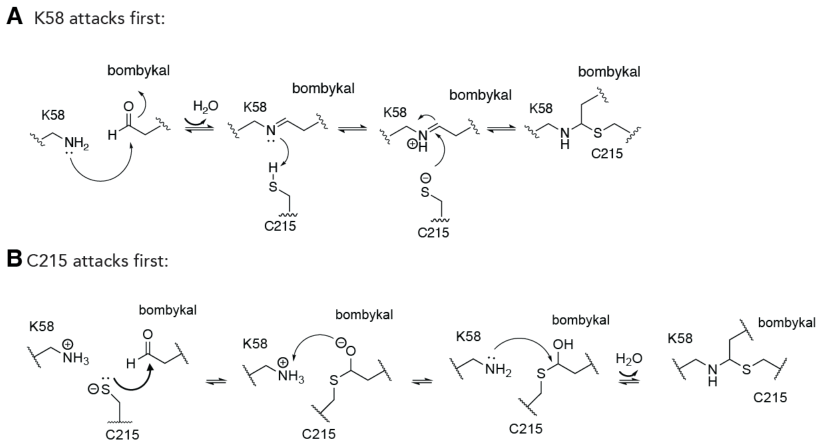


**Figure S5. Proposed mechanisms of thio-hemiaminal formation.** Schematic of the formation of the covalent linkage between the aldehyde functional group of bombykal and residues Lys58 and Cys215. Two possible reaction pathways are shown: **(A)** An initial nucleophilic attack by Lys58 to form a Schiff base intermediate, followed by a secondary attack from the Cys215 thiolate to form the thio-hemiaminal. **(B)** An initial nucleophilic attack by the Cys215 thiolate (present in large fraction due to the proximity to the Lys58 base) followed by deprotonation of Lys58 and subsequent nucleophilic attack to form the thio-hemiaminal.

### Figure S6

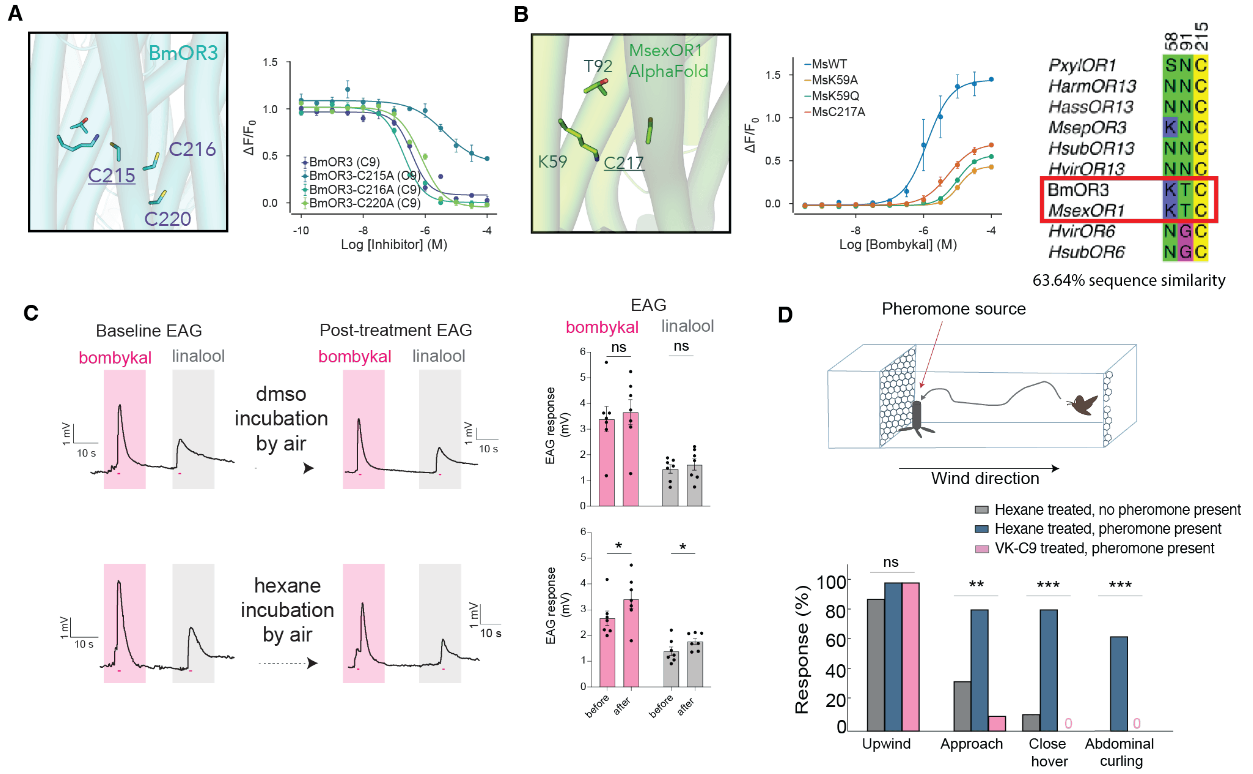


**Figure S6. Control experiments for *in vitro*, *ex vivo* and *in vivo* inhibitor assays. (A)** Left, structural view of all the cysteine residues available in the BmOR3 binding pocket. Right, dose-dependent inhibition of BmOR3 constructs in which individual cysteines in the binding pocket were mutated to an alanine. Activation was measured against a fixed bombykal concentration post incubation in C9-ACA. **(B)** Left, Corresponding view of the predicted MsexOR1 binding pocket (AlphaFold)**.** Middle, Bombykal dose-response curves for MsOR1/MsOrco mutants targeting residues predicted to form a thio-hemiaminal with the aldehyde group of bombykal. Right, sequence alignment showing the conserved residues in the polar end of the pocket. **(C)** Representative EAG traces of *M. sexta* antennae in response to bombykal (magenta) and linalool (grey) before (left) and after (middle) treatment with solvent controls. Peak voltage changes changes for each stimulus are quantified (right). n = 7 antennae per group. Pink lines beneath traces indicate 1-s stimulation. Bars represent mean ± SEM with dots indicating the individual replicates. Asterisks indicate significant difference between groups by two-way repeated measures ANOVA with Sidak's post hoc test $*p < 0.05$. **(D)** Schematic of experimental setup and *M. sexta* flight path in the wind tunnel upwind toward the odor source (top). Percentage of moths pre-exposed with hexane (grey and blue) or C-9 vinyl-ketone (pink) performing non-pheromone behavior (grey) or pheromone-related behaviors (blue and pink). n = 9-11 moths. The differences among each the treatment for each behavior:

upwind flight ($\chi^2 = 2.414$, $p = 0.299$); *approach* ($\chi^2 = 11.526$, ** $p = 0.003$); close hover ($\chi^2 = 18.636$, *** $p < 0.001$); abdomen curling ($\chi^2 = 15.771$, *** $p < 0.001$).

### Table S1

| | BmOR3/BmOrco<br>**Bombykal**<br>(PDB: XXXX)<br>(*EMD-XXXXX)* | BmOR3/BmOrco<br>**Bombykol**<br>(PDB: XXXX)<br>(*EMD-XXXXX)* | BmOR3/BmOrco<br>**Apo**<br>(PDB: XXXX)<br>*(EMD-XXXXX)* |
|---|---|---|---|
| **Data collection and Processing** | | | |
| Magnification | 81,000 | 81,000 | 81,000 |
| Voltage (kV) | 300 | 300 | 300 |
| Pixel size at detector (Å /pixel) | 1.061 | 1.061 | 1.061 |
| Total Electron exposure ($e^-$/Å$^2$) | 50 | 50 | 50 |
| Defocus range (μm) | -1 to -2 | -1 to -2 | -1 to -2 |
| Tilt angle (°) | 30 | 30 | 30 |
| Micrographs used | 6,320 | 9,768 | 5389 |
| Total extracted particles | 4,498,500 | 6,058,000 | 4,652,000 |
| Final particles | 1,070,549 | 1,191,563 | 326,678 |
| Symmetry imposed | C1 | C1 | C1 |
| Map resolution (Å) | 2.68 | 2.81 | 2.89 |
| FSC threshold | 0.143 | 0.143 | 0.143 |
| **Refinement** | | | |
| Map sharpening B factor (Å$^2$) | -111 | -126 | -100.3 |
| Model composition | | | |
| Non-hydrogen atoms | 12,963 | 12,960 | 12,951 |
| Protein residues | 1,612 | 1,612 | 1,612 |
| Ligands | 1 | 1 | 0 |
| R.m.s deviations | | | |
| Bond length (Å) | 0.004 | 0.003 | 0.005 |
| Bond angle (°) | 0.595 | 0.515 | 0.618 |
| Validation | | | |
| MolProbity score | 1.85 | 1.54 | 1.75 |
| MolProbity clash score | 5.39 | 4.19 | 6.66 |

| | | | |
|---|---|---|---|
| Rotamer outliers (%) | 5.39 | 2.77 | 2.91 |
| Ramachandran plot | | | |
| Favored (%) | 98.68 | 99.19 | 97.87 |
| Allowed (%) | 1.32 | 0.81 | 2.13 |
| Disallowed (%) | 0 | 0 | 0 |

**Table S1. Cryo-EM data collection, refinement, and validation statistics**

### Table S2

| Construct | n | Ligand | Baseline (Raw F± SE) | Mean (-log $EC_{50}$ ± SE) | Max (ΔF/F ± SE) | Hill Coefficient (n± SE) |
|---|---|---|---|---|---|---|
| No OR | 12 | - | 6302.6 ± 125.3 | - | - | - |
| BmOR3-WT | 12 | Bombykal | 11274.1± 148.6 | 7.001 ± 0.032 | 1.020 ± 0.013 | 1.29 ± 0.11 |
| BmOR3-K58A | 12 | Bombykal | 10994.3± 132.5 | 5.147 ± 0.046 | 0.189 ± 0.005 | 1.22 ± 0.13 |
| BmOR3-K58Q | 12 | Bombykal | 10781.6± 142.0 | 5.055 ± 0.010 | 0.622 ± 0.005 | 1.66 ± 0.05 |
| BmOR3-C215A | 12 | Bombykal | 9796.8± 154.7 | 5.491 ± 0.028 | 0.935 ± 0.018 | 1.66 ± 0.16 |
| BmOR3-WT | 12 | Bombykol | 11889.6 ± 175.6 | 5.826 ± 0.017 | 0.758 ± 0.007 | 1.28 ± 0.06 |
| BmOR3-K58A | 12 | Bombykol | 11920.6 ± 138.6 | 5.108 ± 0.025 | 0.213 ± 0.004 | 1.43 ± 0.10 |
| BmOR3-K58Q | 12 | Bombykol | 10824.9 ± 134.4 | 5.419 ± 0.025 | 0.828 ± 0.013 | 1.37 ± 0.09 |
| BmOR3-C215A | 12 | Bombykol | 10634.3 ± 153.3 | 5.256 ± 0.018 | 0.702 ± 0.008 | 1.38 ± 0.07 |
| BmOR3-WT | 12 | Bombykal | 13148.6 ± 291.1 | 6.674 ± 0.067 | 1.011 ± 0.027 | 1.09 ± 0.16 |
| BmOR3-F159A | 6 | Bombykal | 8195.2 ± 196.3 | 5.240 ± 0.163 | 0.163 ± 0.015 | 1.30 ± 0.52 |
| BmOR3-L349A | 6 | Bombykal | 6164.7 ± 126.5 | 5.485 ± 0.048 | 0.555 ± 0.018 | 1.62 ± 0.26 |
| BmOR3-V352A | 6 | Bombykal | 10076.8 ± 189.1 | 6.283 ± 0.044 | 0.662 ± 0.016 | 1.80 ± 0.27 |
| BmOR3-V348A | 6 | Bombykal | 11207.4 ± 201.2 | 6.326 ± 0.034 | 1.267 ± 0.020 | 1.16 ± 0.09 |
| BmOR3-L353A | 6 | Bombykal | 10623.3 ± 231.1 | 6.332 ± 0.062 | 0.937 ± 0.026 | 1.06 ± 0.14 |
| MsOR1-WT | 2 | Bombykal | 13055.6 ± 179.1 | 5.892 ± 0.044 | 1.434 ± 0.034 | 1.23 ± 0.13 |
| MsOR1-K59A | 2 | Bombykal | 14031.4± 191.5 | 4.961 ± 0.032 | 0.428 ± 0.012 | 1.89 ± 0.25 |
| MsOR1-K59Q | 2 | Bombykal | 12562.3± 163.9 | 5.006 ± 0.023 | 0.551 ± 0.009 | 1.55 ± 0.11 |
| MsOR1-C217A | 2 | Bombykal | 8884.5± 143.9 | 5.267 ± 0.035 | 0.677 ± 0.013 | 1.17 ± 0.09 |

**Table S2. Wild-type BmOR3/Orco and MsOR1/Orco dose-response parameters.**

**Table S3**

| Construct | n | Inhibitor | Agonist (Name, X μM) | Baseline | Mean (-log $IC_{50}$ ± SE) | Max (ΔF/F ± SE) | Hill Coefficient (n± SE) |
|---|---|---|---|---|---|---|---|
| BmOR3-WT | 3 | C4-ACA | kal, 0.63 | 10975.7± 174.0 | 4.086 ± 0.318 | 0.230 ± 0.030 | - 1.03 ± 0.22 |
| BmOR3-WT | 3 | C5-ACA | kal, 0.63 | 10838.3± 150.1 | 5.219 ± 0.077 | 0.049 ± 0.057 | - 1.46 ± 0.31 |
| BmOR3-WT | 3 | C6-ACA | kal, 0.63 | 11084.6± 163.9 | 5.749 ± 0.115 | - 0.013 ± 0.043 | - 0.78 ± 0.14 |
| BmOR3-WT | 3 | C7-ACA | kal, 0.63 | 11353.3± 274.4 | 6.250 ± 0.077 | - 0.004 ± 0.035 | - 0.95 ± 0.14 |
| BmOR3-WT | 3 | C8-ACA | kal, 0.63 | 10016.4± 160.2 | 6.775 ± 0.048 | 0.022 ± 0.024 | - 1.52 ± 0.22 |
| BmOR3-WT | 3 | C9-ACA | kal, 0.63 | 9822.8± 183.4 | 7.044 ± 0.086 | 0.045 ± 0.023 | - 5.00 ± 9.47 |
| BmOR3-WT | 3 | C9-ACA | kal, 3.98 | 11674.9± 173.5 | 6.435 ± 0.087 | 0.083 ± 0.039 | - 1.52 ± 0.41 |
| BmOR3-C215A | 3 | C9-ACA | kal, 3.98 | 10422.0± 179.2 | 5.378 ± 0.220 | 0.463 ± 0.048 | - 0.82 ± 0.26 |
| BmOR3-C216A | 3 | C9-ACA | kal, 3.98 | 10227.7± 190.0 | 6.688 ± 0.058 | 0.004 ± 0.027 | - 1.36 ± 0.21 |
| BmOR3_C220A | 3 | C9-ACA | kol, 3.98 | 9459.6± 189.1 | 6.129 ± 0.074 | - 0.044 ± 0.038 | - 1.16 ± 0.20 |
| BmOR1_WT | 6 | C4-ACA | kol, 0.63 | 13293.3 ± 160.1 | - | 1.290 ± 0.069 | - |
| BmOR1_WT | 6 | C5-ACA | kol, 0.63 | 12570.3± 156.6 | - | 1.172 ± 0.066 | - |
| BmOR1_WT | 6 | C6-ACA | kol, 0.63 | 12833.3± 148.2 | - | 1.048 ± 0.052 | - |
| BmOR1_WT | 6 | C7-ACA | kol, 0.63 | 12741.1± 199.5 | - | 1.020 ± 0.048 | - |
| BmOR1_WT | 6 | C8-ACA | kol, 0.63 | 11515.5± 169.8 | - | 1.038 ± 0.059 | - |
| BmOR1_WT | 6 | C9-ACA | kol, 0.63 | 11631.6± 204.3 | - | 1.077 ± 0.080 | - |
| MsOR1-WT | 4 | C4-ACA | kal, 1.58 | 12829.3 ± 177.7 | 5.469 ± 0.104 | 0.070 ± 0.065 | - 1.14 ± 0.26 |
| MsOR1-WT | 4 | C5-ACA | kal, 1.58 | 12676.1 ± 183.7 | 5.810 ± 0.157 | - 0.044 ± 0.080 | - 0.87 ± 0.24 |
| MsOR1-WT | 4 | C6-ACA | kal, 1.58 | 12653.7 ± 185.5 | 6.824 ± 0.091 | 0.012 ± 0.050 | - 1.65 ± 0.48 |
| MsOR1-WT | 4 | C7-ACA | kal, 1.58 | 12246.6 ± 224.6 | 7.390. ± 0.050 | 0.007 ± 0.029 | - 2.72 ± 0.79 |
| MsOR1-WT | 4 | C8-ACA | kal, 1.58 | 10990.7 ± 198.6 | 7.698 ± 0.059 | 0.007 ± 0.029 | - 2.60 ± 0.62 |
| MsOR1-WT | 4 | C9-ACA | kal, 1.58 | 11128.1 ± 336.3 | 7.783 ± 0.069 | - 0.009 ± 0.032 | - 2.12 ± 0.54 |

**Table S3. Covalent inhibitor against BmOR3, BmOR1, and MsOR1 dose-response parameters.**

### Table S4

| Compound | Boiling Point, °C | Pressure (Boiling Point), Torr | Boiling Point corrected to 1 atm, ºC | Reference |
|---|---|---|---|---|
| Bombykal | 135 - 145 | 0.01 | 419 - 435 | *Tetrahedron Lett*, **1977**, 121 – 124. |
| Bombykol | 118 - 119 | 0.0004 | NA | *Tetrahedron Lett*, **1989**, 243 – 246. |
| | 128 – 129 | 0.01 | 408 – 410 | *Tetrahedron*, **1984**, 2741 – 2750. |
| | 119 - 120 | 0.01 | 394 – 396 | *Tetrahedron*, **1980**, 1215 – 1222. |
| | 115 – 130 | 0.01 | 388 – 412 | *Tetrahedron Lett*, **1977**, 121 – 124. |
| | 119 – 120 | 0.001 | NA | *Tetrahedron Lett*, **1975**, 1465 – 1468. |
| | 140 – 150 | 0.01 | 427 – 443 | *Liebigs Ann Chem,* **1962**, 39 – 64. |
| | 130 – 133 | 0.005 | NA | *Liebigs Ann Chem*, **1962**, 65 – 90. |
| C4-ACA | 110 | 7 | 251 | *J Am Chem Soc*, **1956**, 2556 – 2557. |
| C5-ACA | 86 - 87.5 | 0.01 | 342.5 – 344.9 | *Helv Chim Acta*, **1972**, 1883 – 1897. |
| | 82 | 0.5 | 271.3 | *J Am Chem Soc*, **1956**, 2556 – 2557. |
| | 78 – 80 | 0.1 | 296.0 - 299.0 | US Patent 2768202, **1955**. |
| C6-ACA | 95 – 105 | 0.2 | 308.6 - 322.9 | US Patent 2768202, **1955**. |
| VK-C9 | 74 | 0.4 | 265 | *Tetrahedron Lett*, **1980**, 789 – 792. |
| | 98 | 2.5 | 259 | *Chem Ber*, **1977**, 1007 – 1019. |

**Table S4. Boiling point data summary on chloroacetamide, vinyl ketone, bombykol, bombykal.**
